# T cell fate is dictated by different antigen presenting cells in response to dietary *versus* gut epithelial self-antigen

**DOI:** 10.1101/2025.11.06.687030

**Authors:** Yixuan D. Zhou, Hailey Brown, Emily Schaffer, Gwen M. Taylor, Kay L. Fiske, Macy R. Komnick, Sebastian Lopez, Terence S. Dermody, Daria Esterházy

## Abstract

We investigated whether T cell responses and antigen-presenting cell (APC) requirements in gut-draining lymph nodes differ by antigen source, diet *versus* epithelium. Using mice fed ovalbumin (OVA) or expressing secreted (s), cytosolic (c), or transmembrane (tm) epithelial OVA, we compared OVA-specific T cell fates. At baseline and after reovirus infection, T cell responses were comparable across models. However, helminth infection induced Th2 cell polarization in sOVA and tmOVA but not cOVA or OVA-fed mice. BATF3⁺ APCs were indispensable for CD4⁺ T cell proliferation only in cOVA mice yet drove Treg cell differentiation across all epithelial OVA models. In contrast, antigen presentation by RORγt⁺MHC-II⁺ APCs was exclusively required for Treg cell induction by dietary OVA. These distinct APC dependencies correlated with susceptibility to pathology elicited by dietary *versus* epithelial self-antigens. Thus, antigen origin and presentation context are integrated to shape T cell fate, a new framework for predicting gut immune outcomes.

## Introduction

The intestinal immune system is constantly exposed to dietary and microbial antigens. Although less studied, it also encounters self-antigens, which are likely presented in abundance due to the epithelium’s large surface area, rapid turnover rate, the absorptive nature of intestinal epithelial cells, and the vast lymphatics. The reaction to dietary antigens is well characterized: Under homeostatic conditions, food proteins elicit peripheral Treg (pTreg) cell induction, leading to their tolerance. In contrast, viral, helminthic, and bacterial infections lead to a Th1, Th2, or often Th17/22 cell response, respectively, required for pathogen clearance^1–5^. The default pTreg cell response to dietary antigens and corresponding oral tolerance can be diminished if food antigens are encountered for the first time during any of these infections^1,3,6^, and this may contribute to diseases like celiac disease (CD) or food allergies. Since antigen presenting cells (APCs) in the gut-draining lymph nodes (gLNs) determine T cell fate, the impact of infections on how food antigens are reacted to reflects a switch in the APC subset presenting these antigens or changes in the APC transcriptional and functional profiles^2,3,6^. Additionally, recent work has shown that while dietary antigen is loaded onto various APC subtypes^2^, including conventional dendritic cells type 1 and 2 (cDC1 and cDC2) and RORγt⁺MHC-II⁺ APCs (recently more commonly referred to as Thetis cells (TCs)), all of which are capable of inducing pTreg cells^7–10^, only TCs are indispensable for generating food antigen-specific pTreg cells^8–10^.

Whether the same principles for Treg induction in the periphery apply to self-antigens is unknown. In fact, T cell responses to another major source of antigens in the gut, the commensal microbiota, are highly strain specific, and while when pTreg cells to commensals are induced this process seems to also depend on TCs^8^, not all commensals induce pTreg cells. For example, segmented filamentous bacteria (SFB) drive Th17 cell differentiation^4^, *Helicobacter hepaticus* promotes RORγt⁺ pTreg cell induction in gLNs^11^, and *Akkermansia muciniphila* elicits a Tfh cell-dominated response, primarily in the Peyer’s Patches^12^. As observed for dietary antigens, these dominant T cell fates can shift when antigen presentation occurs in a different inflammatory or tissue context^11–13^. The differences in the nature of the T cell response to dietary and microbial antigens or between microbial strains can only to some extent be explained by the different locations along the gastrointestinal tract at which the antigens are encountered^1^, as homeostatic T cell responses can differ at the same location. Thus, these different responses likely also reflect the selective access of APC subsets to specific antigens and adjuvants that come with these antigens, shaping the functional profiles of the APCs. It is therefore not evident what the rules governing self-reactive T cell fate would be. While central tolerance eliminates most self-reactive T cell clones or redirects them into the Treg cell lineage, additional peripheral tolerance mechanisms are required to prevent autoimmunity. Indeed, studies using mice expressing cytosolic OVA have found that OVA-specific CD8+ T cells develop and home to the gut but acquire a CD8+ Treg cell phenotype rather than destroying the epithelium^14,15^. On the other hand, conditions like gluten-evoked autoimmunity in CD, autoantibodies in inflammatory bowel disease, and the destruction of gut epithelium in patients treated with immune checkpoint inhibitors illustrate the consequences of failed tolerance to intestinal self-antigens^3,16,17^. Thus, there is a need to better understand what determines T cell fate to gut self-antigens, which include a large repertoire of secreted, membrane-bound, and cytosolic proteins.

In this study, we asked whether T cell responses and APC requirements differ between epithelial self- and diet-derived antigens, and among epithelial antigens, whether the subcellular compartment of antigen origin influences immune outcomes. We hypothesized that any such distinctions would result in differential susceptibility to collateral damage following infections or the loss of specific APC subtypes. More broadly, our goal was to uncover principles linking antigen source and presentation context to T cell fate in the gut.

## Results

### Intestinal epithelial self-antigens and oral antigen induce similar antigen-specific FOXP3+ T cell levels in an antigen dose-dependent manner

To directly compare T cell fate and APC requirements following exposure to dietary and self-antigens without confounding factors such as differential T cell receptor (TCR) affinity, we engineered mice that express the dietary antigen ovalbumin (OVA) as a self-antigen. We cloned full-length secreted OVA (sOVA) after a loxP-flanked STOP cassette, allowing tissue-specific Cre recombinase to excise the cassette and place *OVA* under the control of the artificial CAG promoter^18^. We inserted the construct into the *Rosa26* locus using CRISPR (Fig. 1A). We also engineered a version that lacked the internal secretion signal, rendering OVA cytosolic (cOVA), as well as a version lacking the signal peptide but N-terminally fused to the transferrin receptor (TFR) transmembrane domain (tmOVA)^14,15,19^. Each construct included a C-terminal 3×FLAG tag for protein detection purposes. Correct targeting to the mouse genome was confirmed by PCR (Fig. S1A) and sequencing. Mice with these *OVA* transgenes did not stimulate the proliferation of adoptively transferred, OVA-specific CD4+ T cells (OT-II cells) in the gLNs (Fig. S1B) and did not express *OVA* in any tissue tested (Fig. S1C-E). However, following interbreeding with *Villin^CRE-ERT^*^2^ mice^20^, in which *Cre* is expressed in intestinal epithelial cells and translocates to the nucleus after tamoxifen administration, all three *OVA* mouse lines robustly expressed *OVA* along the gut. The expression levels were comparable between the three lines, albeit trended to be lower in *Villin^cOVA^* and *Villin^tmOVA^* than *Villin^sOVA^* mice (Fig. S1F, G). The expected protein sizes were confirmed by immunoblotting, with higher OVA abundance in the jejunal lysates from *Villin^sOVA^* mice than the other two lines (Fig. S1H), potentially because the latter two are less stable, though immunofluorescence staining suggested comparable epithelial expression - and expected localization - across all lines (Fig. S1I).

**Figure 1.**
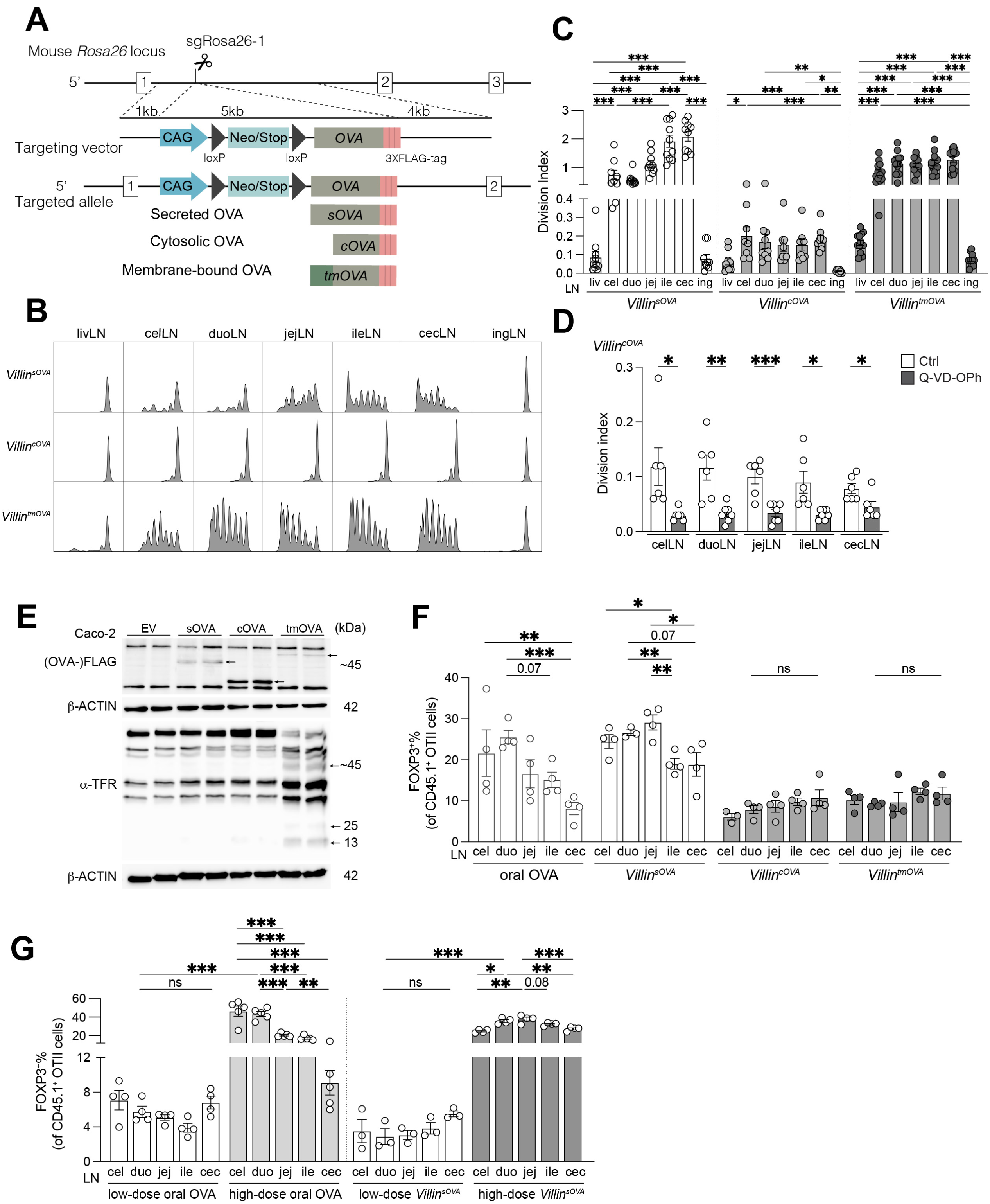
Generation and characterization of mice expressing OVA in different subcellular epithelial cell compartments. **A**. Scheme of transgenic mouse generation. Targeted allele with the indicated transgene was inserted into the *Rosa26* locus using CRISPR/Cas9. **B-C**. CFSE dilution flow plots (**B**) and division index (**C**) of OT-II cells in the indicated LNs 96 h after transfer into *Villin^sOVA^*, *Villin^cOVA^* or *Villin^tmOVA^* mice (*n* = 10-16 per group). **D**. Division index of OT-II cells 96 h after transfer into *Villin^cOVA^*mice treated with pan-caspase inhibitor Q-VD-OPh (*n* = 6 per group). **E**. Western blot of cell lysates of Caco-2 cells transduced with lentivirus encoding empty vector, *sOVA*, *cOVA* or *tmOVA* blotted using antibodies against FLAG, α-TFR and β-ACTIN. Arrows indicate OVA signals. **F**. Frequencies of FOXP3^+^ among total OT-II cells in the indicated LNs 96 h after transfer into mice fed OVA or *Villin^sOVA^*, *Villin^cOVA^* and *Villin^tmOVA^* mice (*n* = 4 per group, as indicated by symbols). **G**. Frequencies of FOXP3^+^ among total OT-II cells in the indicated LNs 96 h after transfer into mice fed high- or low-dose OVA or mice with high or low expression of sOVA (*n* = 4-5 per group, as indicated by symbols). Data are pooled from two or three independent experiments in C, D. Data are representative of two independent experiments in F, G. * p<0.05, ** p<0.01, *** p<0.001 by one-way ANOVA and t-test.

To assess whether the subcellular origin of epithelial antigen influences T cell stimulation, we adoptively transferred CFSE loaded OT-II cells into *Villin^sOVA^*, *Villin^cOVA^*, and *Villin^tmOVA^* mice and quantified their proliferation in gLNs as an activation proxy. In all three mouse lines, OT-II cells proliferated in the gLNs but not the non-draining liver and inguinal LNs (Fig. 1B, C). However, while OT-II cell division was comparable between *Villin^sOVA^* and *Villin^tmOVA^*mice, it was much lower in *Villin^cOVA^* mice. We thought it was unlikely that this result was due to the slightly lower OVA expression in these mice (Fig. S1F-H). Rather, we hypothesized that accessing cytosolic OVA requires epithelial turn over. Consistent with this hypothesis, pharmacological inhibition of apoptosis using a pan-caspase inhibitor (Fig. S2A, B) in *Villin^cOVA^*mice almost completely abrogated OT-II proliferation (Fig. 1D, Fig. S2C). However, proliferation in *Villin^sOVA^* mice was unaffected and only partially reduced in *Villin^tmOVA^* mice by caspase inhibition (Fig. S2D, E), suggesting that antigen access was either independent or only partially dependent on cell death, respectively, in these two mouse lines. As OT-II proliferation in *Villin^tmOVA^* mice displayed an intermediate phenotype compared to *Villin^sOVA^* and *Villin^cOVA^* mice, we tested whether tmOVA also is released from epithelial cells. One potential mechanism is exosomal secretion of transmembrane proteins. However, treating mice with exosome inhibitor GW4869^21^ had no impact on OT-II division (Fig. S2F), arguing against this pathway. Ectodomain shedding is an alternative mechanism that does not require cell death. The tmOVA construct used in our experiments, based on published tmOVA constructs^14,15,19^, retains the ADAM protease cleavage site (Arg100)^22^ within the TFR transmembrane stalk (amino acids 1-116 of the full-length 760-aa human TFR). To test whether ectodomain shedding is the responsible mechanism, we overexpressed sOVA, cOVA, or tmOVA in Caco-2 or HEK293T cells and probed the cell lysates with an antibody specific for the N-terminal domain of TFR^22^. In tmOVA-expressing cells, TFR antibody detected the expected ∼50 kDa full-length tmOVA protein as well as two lower-molecular weight species, one at ∼13 kDa, the size of the N-terminal cleavage product, and another at ∼25 kDa likely representing its dimer^22^ (Fig. 1E, Fig. S2G). However, in the supernatant of HEK293T cells, much less OVA shed from tmOVA was present relative to sOVA (Fig. S2H), indicating that while tmOVA can be shed, sOVA remains the more abundant soluble form.

We next addressed if the subcellular compartment of epithelial OVA influences CD4+ T cell fate at homeostasis and how it compares with the response to dietary OVA. Naïve OT-II cells were simultaneously transferred into C57Bl/6 mice fed OVA via oral gavage and drinking water and into *Villin^sOVA^*, *Villin^cOVA^*, and *Villin^tmOVA^* mice. Cells in the gLNs were analyzed four days later for the expression of activation, lineage and anergy markers: CD44 (early activation), FOXP3 (pTreg), TBET (Th1), GATA3 (Th2), RORγt (Th17), PD-1+CXCR5+BCL6 (Tfh), and FR4+CD73 (anergy). Nearly all OT-II cells acquired CD44, indicating that they had encountered cognate antigen (*data not shown*). Of all lineages, only FOXP3^+^ cells were recovered among OT-II cells at percentages above 2-4%, and the OT-II cells failing to upregulate any lineage-defining transcription factors adopted an anergic phenotype (Fig. 1F, Fig. S2I, and *data not shown*). Oral OVA and sOVA elicited similar percentages of FOXP3+ OT-II cells, with higher induction in small-intestine relative to large-intestine draining LNs (Fig. 1F), as previously reported^1^. By contrast, cOVA and tmOVA induced only about half the amount of FOXP3+ cells (10% of OT-II cells), and this induction was the same across all gLNs. We reasoned that this reduced and regionally uniform pTreg cell induction in cOVA and tmOVA mice relative to OVA-fed and sOVA mice could reflect differences in antigen dose rather than subcellular compartment. To test this hypothesis, we matched the antigen levels by using a dose of tamoxifen in *Villin^sOVA^* mice that resulted in similar gut OVA protein levels across the OVA lines (Fig. S2J) and comparable OT-II cell division following transfer as in *Villin^cOVA^* mice (compare Fig. S2K and Fig. 1C). We also fed less OVA, which reduced OT-II cell division to *Villin^cOVA^*-like levels (compare Fig. S2K and Fig. 1C). Under these conditions, pTreg cell induction dropped to ∼4-8% of OT-II cells, comparable to *Villin^cOVA^*and *Villin^tmOVA^* mice, and was uniform across all gLNs (Fig. 1C, G).

In sum, we have generated mice that permit the expression of OVA in different subcellular compartments under the control of tissue-specific Cre recombinase of choice. In the context of gut epithelium, the compartment of origin influences antigen availability at steady state, but when antigen dose is matched, epithelial self-antigens and dietary antigen drive comparable pTreg cell differentiation.

### Antigen presentation by cDC1s, but not Thetis cells, is required for pTreg cell induction to gut self-antigens

Two APC subtypes have been implicated as primarily responsible for pTreg cell induction in response to oral antigens: TCs, whose ablation or loss of MHC-II expression abolishes pTreg cell generation^8–10^, and cDC1s, whose loss leads to a 50% reduction of pTreg cells^2,7^ (Fig. S3A). To assess whether the same APC requirements apply to the differentiation of pTreg cells recognizing self-antigens, we interbred *Batf3*^-/-^ mice, which are devoid of cDC1s^23^, with our *Villin^OVA^* lines and generated bone marrow chimeras (BMCs) using *MHCII^11Rorc^* marrow, which lack MHC-II expression on TCs. We confirmed that only the targeted APC was indeed absent in the gLNs of these mice (Fig. S3B-E). Ablation of cDC1 completely prevented OT-II proliferation and expansion in *Villin^cOVA^*mice, but only partially impaired proliferation and expansion in *Villin^sOVA^*and *Villin^tmOVA^* mice (Fig. 2A-C, and *data not shown*). FOXP3+ OT-II cells were reduced by 50-75% in high-dose tamoxifen-treated *Villin^sOVA^* and *Villin^tmOVA^* mice and were almost absent in low-dose tamoxifen-treated *Villin^sOVA^*mice (Fig. 2D-F). As anticipated, the small fraction of T-BET+ cells observed at homeostasis was reduced to a degree similar to FOXP3+ cells in both *Villin^sOVA^* and *Villin^tmOVA^* mice (Fig. 2G, H). In contrast, the opposite picture of dependencies was found when antigen presentation by TCs was impaired. Using the same bone marrow donors, we confirmed that pTreg cell induction following oral OVA administration was blunted in all gLNs, irrespective of antigen dose (Fig. 2I, J). However, TCs only marginally contributed to pTreg cell differentiation in high-dose tamoxifen-treated *Villin^sOVA^* mice, and the loss of antigen presentation by TCs was associated with an increase in pTreg cells in *Villin^cOVA^*, *Villin^tmOVA^* and low-dose tamoxifen-treated *Villin^sOVA^*mice (Fig. 2K-N). In each case, total OT-II numbers and proliferation correlated inversely with pTreg cell frequency. OT-II numbers and proliferation increased following oral OVA and in high-dose tamoxifen-treated *Villin^sOVA^* mice but were relatively unaffected in low-dose tamoxifen-treated *Villin^sOVA^* mice, *Villin^cOVA^*, and *Villin^tmOVA^* mice (Fig. S3F-K).

**Figure 2.**
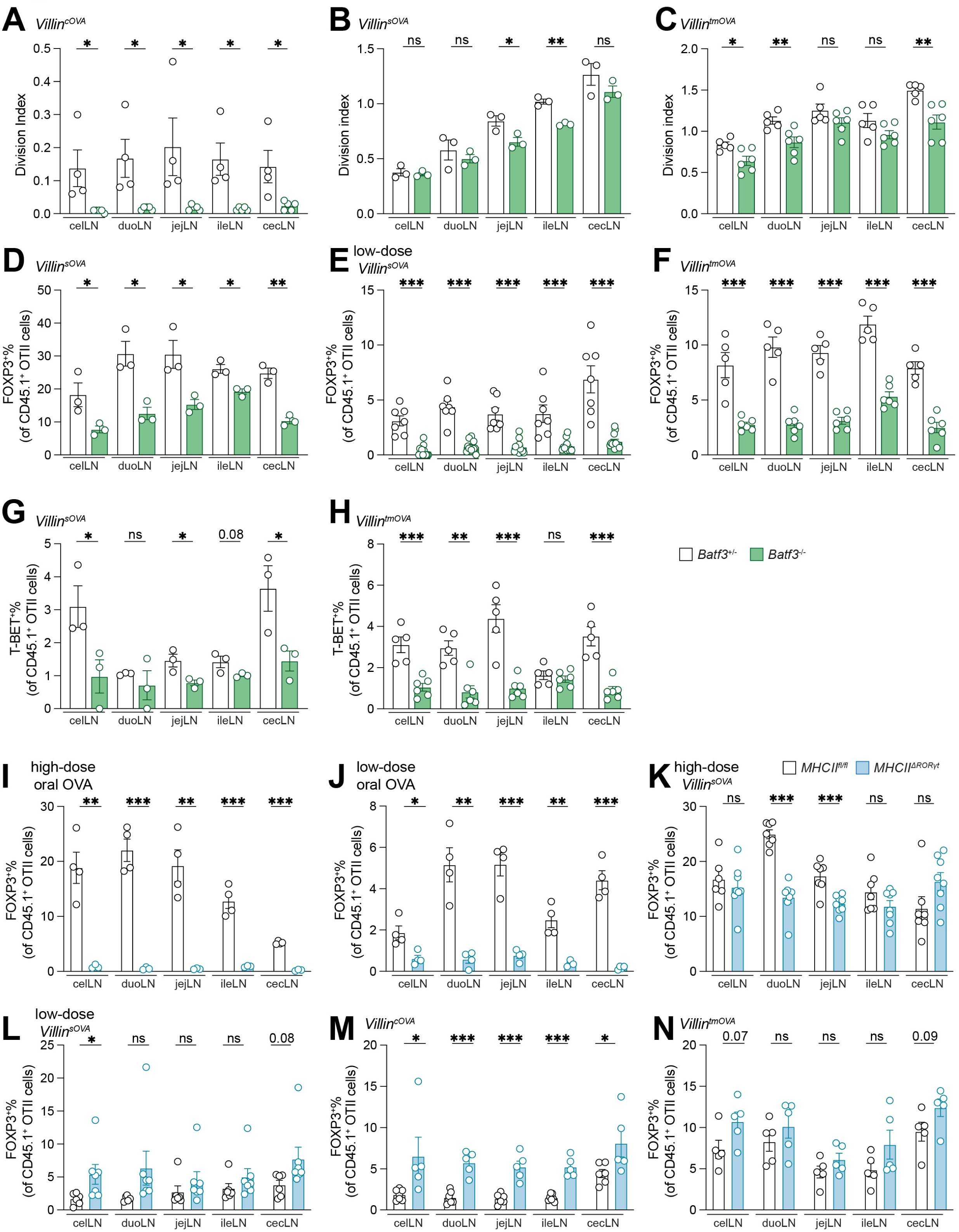
Antigen presentation by cDC1s is required while that by Thetis cells is dispensable for the generation of gut self-antigen specific FOXP3+ T cells. A-C. Division index of OT-II cells 96 h after transfer into *Villin^cOVA^* (**A**), *Villin^sOVA^* (**B**) and *Villin^tmOVA^* (**C**) mice that are *Batf3*-proficient or -deficient. **D-F**. Frequencies of FOXP3^+^ among total OT-II cells in the indicated LNs 96 h after transfer into *Batf3*-proficient or -deficient mice expressing high-dose sOVA (**D**), low-dose sOVA (**E**) or tmOVA (**F**) in the epithelium. **G-H**. Frequencies of T-BET^+^ among total OT-II cells in the indicated LNs 96 h after transfer into *Batf3*-proficient or -deficient mice expressing sOVA (**G**) or tmOVA (**H**) in the epithelium. **I-N**. Frequencies of FOXP3^+^ among total OT-II cells in the indicated LNs 96 h after transfer into BMC receiving *MHCII^fl/fl^*or *MHCII^11RORγt^* BM fed high-dose (**I**) or low-dose OVA (**J**), or mice expressing high-dose sOVA (**K**), low-dose sOVA (**L**), cOVA (**M**) and tmOVA (**N**) in the epithelium. Data are representative of two independent experiments in A-H (*n* = 3-7 per group, as indicated by symbols). Data represent one experiment in I-N (*n* = 4-6 per group, as indicated by symbols). * p<0.05, ** p<0.01, *** p<0.001 by t-test.

These data suggest that presentation of cytosolic antigen is strictly dependent on cDC1s, while secreted antigens are accessed by additional APC subsets. Moreover, unlike luminal, foreign antigens, which rely primarily on presentation by TCs for pTreg cell induction, epithelial self-antigens induce pTreg cells predominantly via cDC1s, with TCs contributing only under conditions of high antigen abundance.

### Secreted but not cytosolic self-antigen induces Th2 cells in response to helminth infection in a cDC2-dependent manner

We next speculated that secreted antigen, like oral antigen^2^, would be presented by the other major APCs in gLNs, cDC2s. cDC2s have several subtypes^24,25^. In the gut, *Klf4*-dependent cDC2Bs are required for the induction of Th2 cells in response to allergens and helminths^26^, while *Notch2*-dependent cDC2As respond to bacteria and fungi, inducing Th17/22 cells^27^. Thus, GATA3 and RORγt induction in CD4 T cells mark cDC2B- and cDC2A-mediated presentation, respectively. We therefore inoculated mice with *S. venezuelensis*, a strong Th2-inducing helminth, and examined OT-II fate in *Villin^OVA^* lines. Notably, dietary OVA does not elicit an interaction between Th2-inducing cDC2s and T cells^2^. However, when OVA came from the epithelium, OT-II cells expanded (Fig. S4A) and over 20% of transferred cells became GATA3+ in the gLNs draining the upper small intestine, the site of helminth infection, in *Villin^sOVA^* and *Villin^tmOVA^* mice (Fig. 3A, B). Analogous to the response to oral antigen, FOXP3+ OT-II cells were diminished in those LNs (Fig. 3A, C), likely due to the relative exclusion of cDC1s from the gut in the context of *S. venezuelensis*^2^. As expected, OT-II cells in *Villin^cOVA^* mice remained unaffected (Fig. 3A-C, Fig. S4A), consistent with the conclusion that presentation of cytosolic self-antigen is primarily dependent on cDC1s. We next tested whether secreted but not cytosolic self-antigens are capable of inducing RORγt+ CD4 T cells, a fate endowed by cDC2As and observed in the context of luminally-sourced OVA^1^. Following infection with *C. rodentium*, a colonic pathogen, the frequency of RORγt+ OT-II cells increased and that of FOXP3+ decreased in the ceco-colonic LN of *Villin^sOVA^*, mirroring responses to oral OVA, but not *Villin^cOVA^* mice (Fig. S4B, C). To further validate that secreted but not cytosolic self-OVA is presented by cDC2s, we ablated this lineage using *Zeb2* enhancer knockout (*Zeb2^111+2+3^*) mice^2,28^, in which cDC2s are reduced by over 80% (Fig. S3B, D). The loss of cDC2s did not impact the low levels of GATA3+ OT-II cells in the gLNs at homeostasis (Fig. S4D, E). However, during *S. venezuelensis* infection, *Zeb2^111+2+3^* mice interbred with *Villin^sOVA^* and *Villin^tmOVA^* mice displayed a blunted OVA-specific Th2 response (Fig. 3D). The blunted Th2 response also was associated with restored pTreg cell induction (Fig. S4F), indicating that cDC1s remain competent to drive tolerance when antigen access is preserved, even during helminth infection.

**Figure 3.**
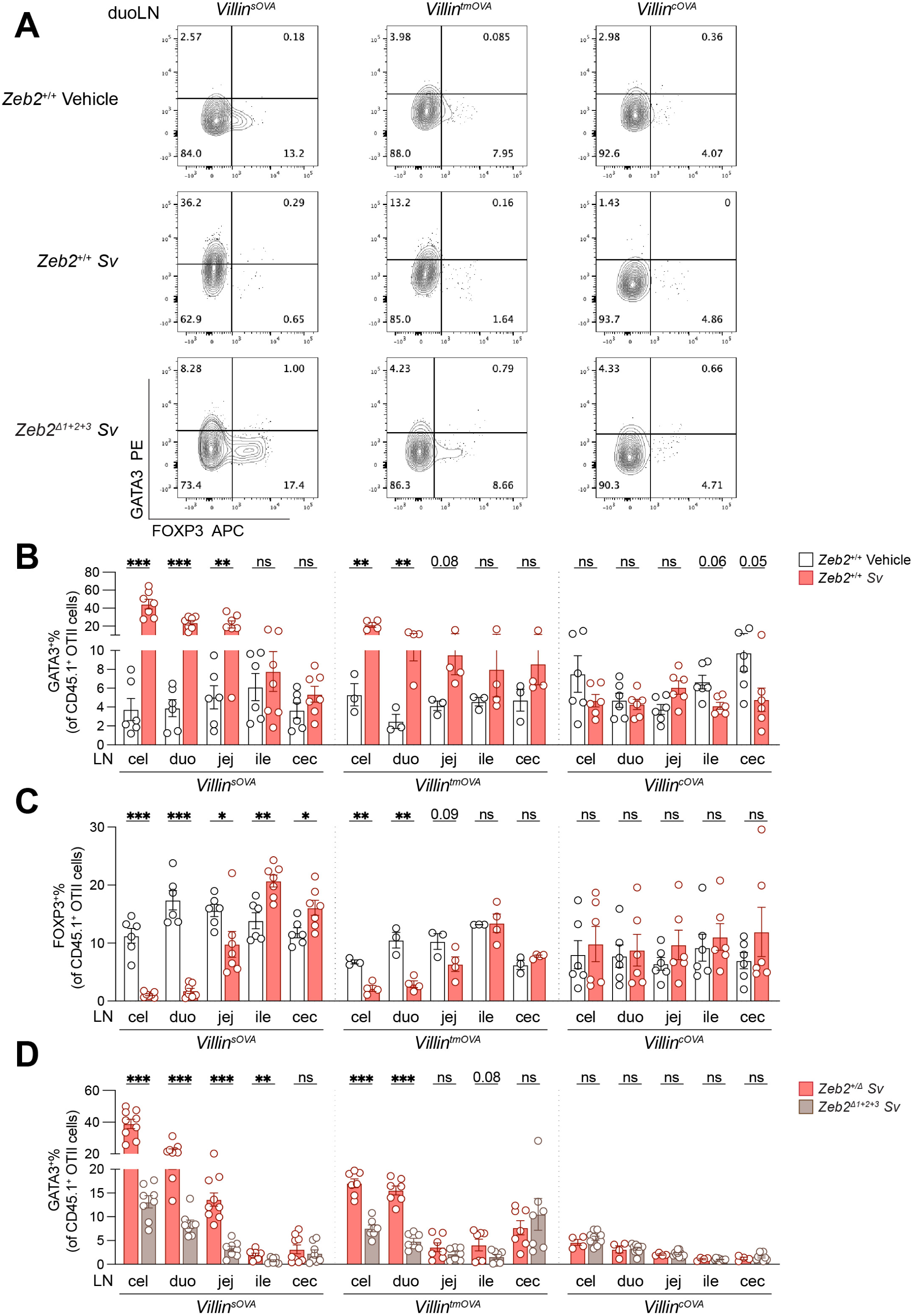
Secreted but not cytosolic self-antigens induce Th2 cells in response to helminth infection in a cDC2-dependent manner. **A.** Flow plots of OT-II cells in duodenal LNs of OVA-expressing mice with the indicated genotypes and *S. venezuelensis* infection status. **B-C.** Frequencies of GATA3^+^ (**B**) or FOXP3^+^ (**C**) among total OT-II cells 72 h after transfer into *Villin^sOVA^*, *Villin^tmOVA^* and *Villin^cOVA^* mice that were *S. venezuelensis*- or vehicle-infected (*n* = 3-7 per group, as indicated by symbols). **D.** Frequencies of GATA3^+^ among total OT-II cells 72 h after transfer into *S. venezuelensis*-infected *Villin^sOVA^*, *Villin^tmOVA^* and *Villin^cOVA^* mice that were deficient for *Zeb2* enhancer and their littermate controls (*n* = 5-9 per group, as indicated by symbols). Data are pooled from two independent experiments in B-D. * p<0.05, ** p<0.01, *** p<0.001 by t-test.

Collectively, these findings indicate that secreted but not cytosolic self-antigen from the gut epithelium is presented by cDC2 subsets that can polarize CD4 T cells towards either GATA3+ and RORγt+ fates following pathogen infection. This response contrasts to that following introduction of dietary antigen, which fails to elicit GATA3+ T cells in the context of helminth infection^1,2^ but is capable of inducing RORγt+ cells.

### Intestinal viral infection similarly alters epithelial self- and oral antigen-specific T cell fate

We next queried the consequences of distinct APC dependencies of oral versus epithelial self-antigens in pathophysiological settings. We predicted that adverse immune outcomes dependent on cDC1s would be similar when stimulated by oral OVA or any *Villin^OVA^* derived antigen. In contrast, immune outcomes of processes dependent on TCs or cDC2s would differ depending on whether antigens are dietary, secreted, or cytosolic. Since cDC1s are critical for inducing antiviral Th1 cells and cytotoxic CD8+ T (T_cyt_) cells, we first tested if infection with reovirus T1L^3,29,30^ would induce ectopic OVA-specific T-BET+ Th1 and T_cyt_ cells and concomitantly reduce FOXP3+ Treg cells equivalently when the antigen was dietary and gut epithelium derived.

Indeed, following OT-II transfer into mice fed OVA or all three *Villin^OVA^* lines, T1L infection induced comparable percentages of T-BET+ OT-II cells and reduced FOXP3+ OT-II cells across all gLNs relative to vehicle controls (Fig. 4A-C, Fig. S5A). Using *Villin^sOVA^* mice, we verified that the observed T-BET induction was dependent on cDC1s (Fig. S5B). Similarly, naïve OVA-specific CD8+ T (OT-I) cells became cytotoxic in the gLNs of all groups after T1L infection, as assessed by the robust emergence of granzyme B (GzmB)+ and interferon γ (IFNγ)+ OT-I cells (Fig. 4D, Fig. S5C-E). An increase in OVA-specific T_cyt_ and Th1 cells, accompanied by a decrease in Treg cells, was detected within the endogenous repertoire using OVA tetramers (Fig. 4E-H, Fig. S5F, G), indicating that this response also occurred in the endogenous T cell pool.

**Figure 4.**
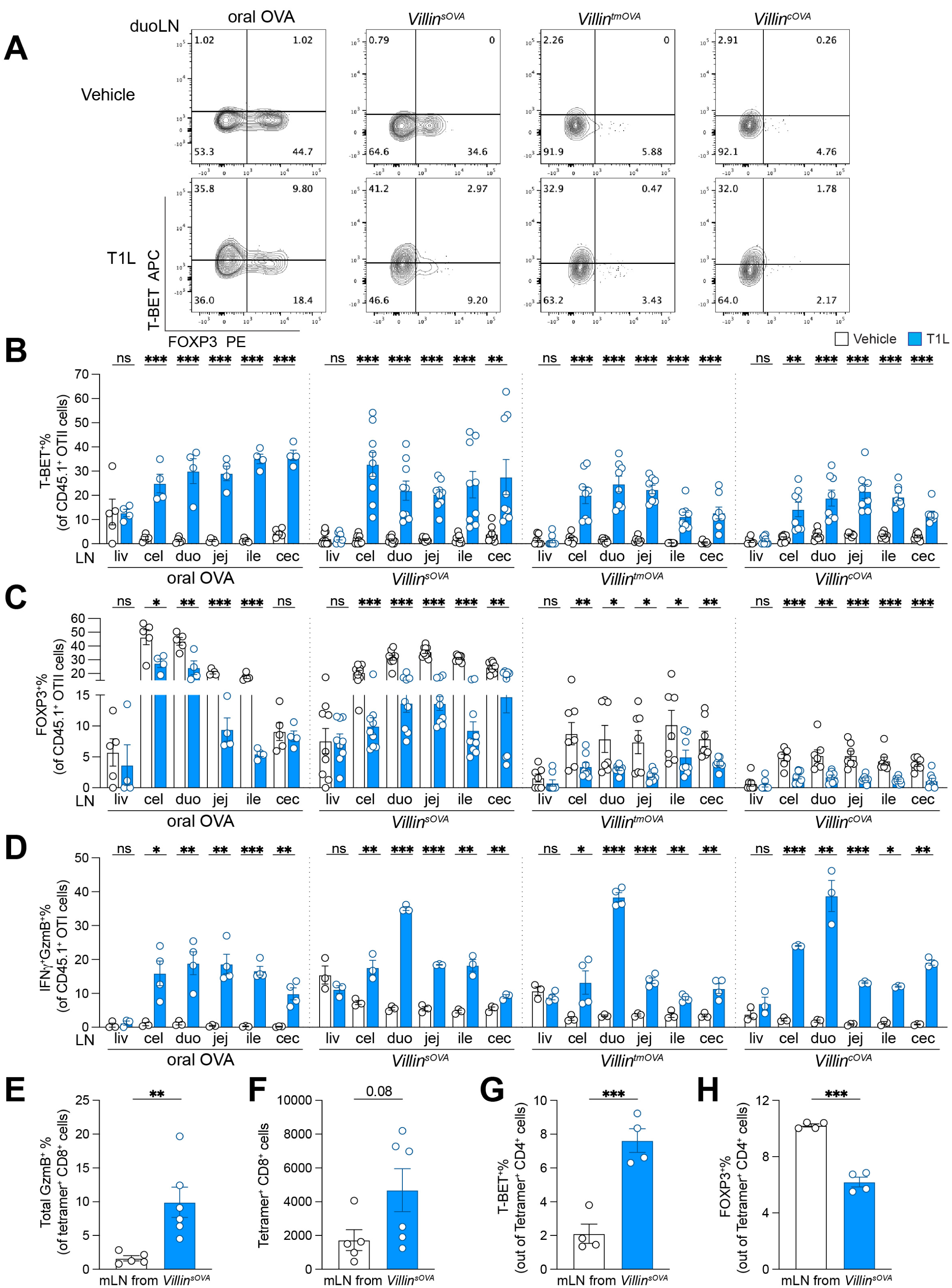
Intestinal viral infection impacts epithelial self-antigen and oral antigen-specific T cell fate similarly. **A.** Flow plots of OT-II cells in duodenal LNs of T1L- or vehicle-infected mice that were fed OVA or express sOVA, tmOVA or cOVA in the epithelium. **B-C.** Frequencies of T-BET^+^ (**B**) or FOXP3^+^(**C**) among total OT-II cells in the indicated LNs 72 h after transfer into T1L- or vehicle-infected mice that were fed OVA or express sOVA, tmOVA or cOVA in the epithelium (*n* = 4-9 per group, as indicated by symbols). **D.** Frequencies of cytotoxic cells among total OT-I cells in the indicated LNs 72 h after transfer into T1L- or vehicle-infected mice that were fed oral OVA or express sOVA, tmOVA or cOVA in the epithelium (*n* = 3-4 per group, as indicated by symbols). **E-F.** Frequencies of GzmB^+^ among total OVA-tetramer^+^CD8^+^ T cells (**E**) and total numbers of OVA-tetramer^+^ CD8^+^T cells (**F**) in the mLNs of T1L- or vehicle-infected *Villin^sOVA^* mice (*n* = 5-6 per group, as indicated by symbols). **G-H.** Frequencies of T-BET^+^ (**G**) and FOXP3^+^ (**H**) among total OVA-tetramer^+^ CD4^+^T cells in the mLNs of T1L- or vehicle-infected *Villin^sOVA^* mice (*n* = 4 per group, as indicated by symbols). Data are pooled from two independent experiments in B, C. Data represent one experiment in D-H. * p<0.05, ** p<0.01, *** p<0.001 by t-test.

Thus, gastrointestinal viral infection leads to similar pro-inflammatory type 1 T cell responses to oral and gut-self antigens in the gLNs. This finding also confirms that oral and self-antigens from any compartment are efficiently presented by cDC1s.

### Intestinal viral infection drives pro-inflammatory T cell infiltration of the gut and comparable tissue damage upon oral and self-antigen recognition

We next determined whether the adverse T cell responses to dietary and self-OVA observed in gLNs following T1L infection translated into the T cell landscape in gut tissue, and if so, whether these responses contributed to gut pathology. OT-I and OT-II cells were adoptively transferred at the time of T1L inoculation, and their abundance and phenotype in the small and large intestine analyzed 7 days later. We used *Villin^sOVA^* mice as a representative for self-antigen, while dietary antigen recognition was modeled by continuous OVA feeding. In both cases, T1L infection led to a higher abundance of OT-I and OT-II cells (Fig. S6A-B), increased GzmB+ OT-I cells and T-BET+ OT-II cells, and reduced FOXP3+ cells in T1L-infected mice than in vehicle controls (Fig. 5A-C, Fig. S6C). Gut tissue destruction by T cells requires both effector T cell infiltration and local inflammatory cues. T1L alone does not lead to sufficient local inflammation to elicit overt pathology^3,31^, likely because viral clearance precedes the return of effector T cells to the gut. However, in the presence of local IL-15, T1L infection can elicit gluten-specific T cells to mediate tissue destruction^3,32^. To test whether T1L-induced OVA-specific Th1 and T_cyt_ cells similarly exacerbate tissue pathology, we administrated indomethacin, a non-steroidal anti-inflammatory drug that induces local gut injury^33^ to mice fed OVA, *Villin^sOVA^* or *Villin^cOVA^* mice 10 days after T1L inoculation and OVA-specific T cell transfer (Fig. 5D). Indomethacin alone caused few mucosal lesions, regardless of whether the transferred T cells encountered cognate antigen or not (Fig. 5E, F). However, if the mice had been inoculated with T1L 10 days prior to indomethacin commencement, the gut of infected mice displayed two to three times as many lesions and was shortened (Fig. 5E-H), with the effects slightly stronger in OVA-fed and *Villin^sOVA^* than *Villin^cOVA^* mice, correlating with the lower antigen dose and slightly lower number of cells induced in this latter model (Fig. 4A-D, Fig. S5A, E). The exacerbated gut pathology occurred in an antigen-specific manner, since T1L failed to aggravate indomethacin-induced pathology in the absence of OVA, even after transfer of OVA-specific T cells (Fig. 5E-H, no OVA group). These results demonstrate that the comparable induction of antigen-specific Th1 and T_cyt_ cells upon oral versus self-antigen in the gLNs translate into similar, antigen-driven gut pathology.

**Figure 5.**
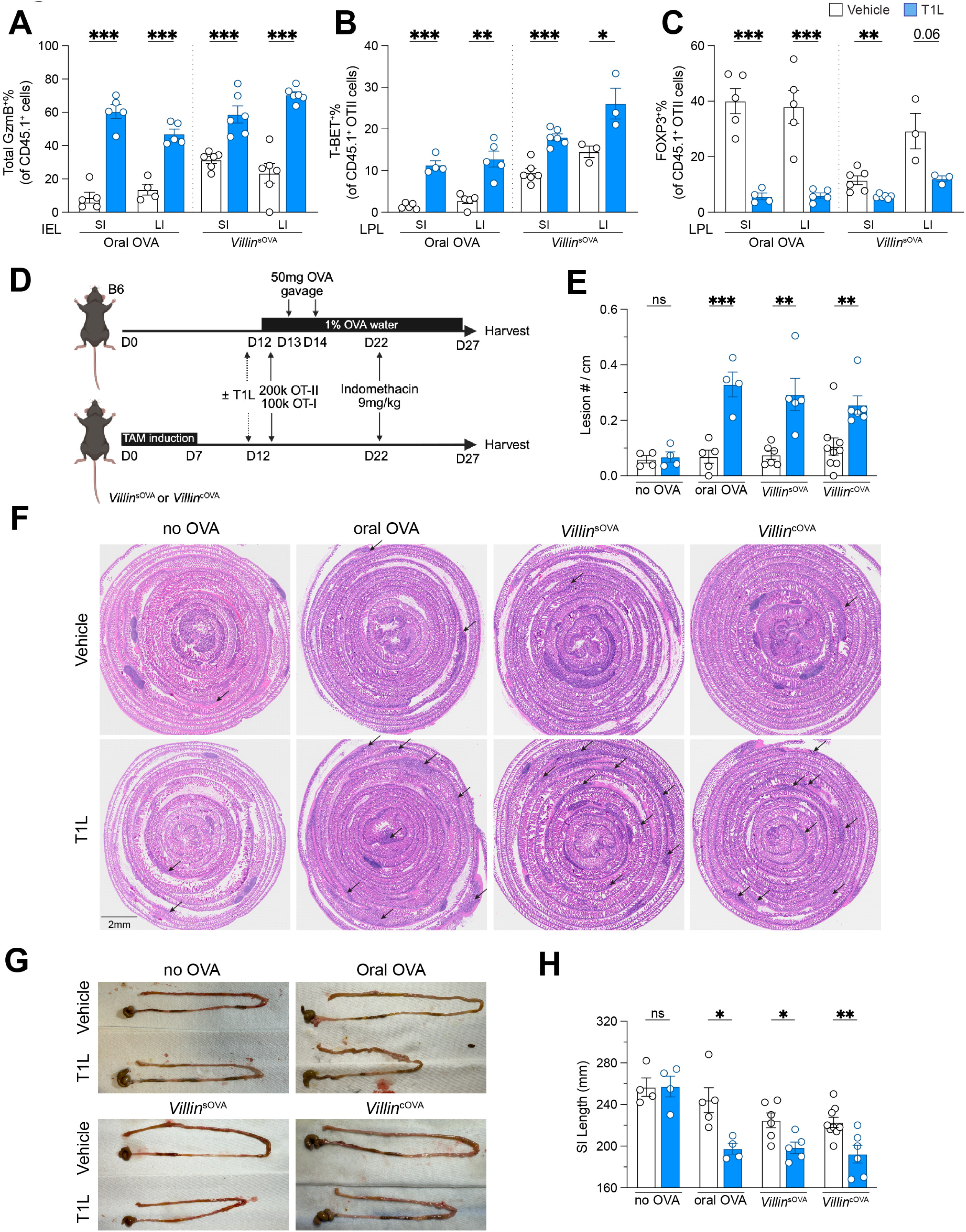
Intestinal viral infection leads to comparable pro-inflammatory T cell infiltration and tissue damage following oral and self-antigen recognition. **A-C.** Frequencies of GzmB^+^ among total OT-I cells (**A**) in the IEL fraction or T-BET^+^ (**B**) or FOXP3^+^ (**C**) among total OT-II cells in the LPL fraction of the gut tissue of T1L- or vehicle-infected mice that were either fed OVA or express sOVA in the epithelium (*n* = 3-6 per group, as indicated by symbols). **D**. Experimental scheme of small intestine damage model. **E**. Lesion number per cm of gut length in each indicated treatment group (*n* = 4-9 per group, as indicated by symbols). **F**. Representative H&E staining of small intestine Swiss roll from each indicated treatment group. The scale bar represents 2 mm. **G**. Representative pictures of small intestines from each indicated treatment group. **H**. Small intestine length of each indicated treatment group (*n* = 4-9 per group, as indicated by symbols). Data are pooled from two independent experiments in A-C. Data represent one experiment in E, H. * p<0.05, ** p<0.01, *** p<0.001 by t-test.

### Antigen presentation by Thetis cells is dispensable for gut self- but not dietary antigen-induced systemic tolerance

Induction of pTreg cells recognizing oral antigens establishes systemic suppression of pro-inflammatory responses to the same antigen, a phenomenon described as oral tolerance. Antigen presentation by TCs, being indispensable for the differentiation of oral antigen-specific pTreg cells, is required for oral tolerance^8–10^. By contrast, we hypothesized that systemic tolerance to gut self-antigens should be independent of TCs, since pTreg cell induction by epithelium-derived OVA was preserved in their absence (Fig. 2K-N). To compare the requirement for TCs in oral *versus* self-antigen induced systemic tolerance, we generated BMCs using *MHCII^11Rorgt^* or *MHCII^fl/fl^*(WT) bone marrow transferred into either *Villin^sOVA^* mice that were later tamoxifen-treated, or C57Bl/6 recipients that were later administered oral OVA. All mice were then immunized with Alum-OVA and challenged with intranasal OVA. We quantified lung immune cell infiltrates and serum anti-OVA IgE and IgG1 levels (Fig. 6A). OVA feeding in the absence of antigen presentation by TCs led to increased immune cell infiltrates in the bronchioalveolar lavage fluid (BALF) and lungs and elevated anti-OVA IgE and IgG1 serum concentrations compared WT BM recipients (Fig. 6B-H, oral OVA groups), as reported previously^8,10^. In contrast, these parameters remained low in *Villin^sOVA^* recipients of *MHCII^11Rorgt^* BM and indistinguishable from WT BM recipients (Fig. 6B-H, *Villin^sOVA^* group), demonstrating that systemic suppression of allergic anti-OVA responses remained intact.

**Figure 6.**
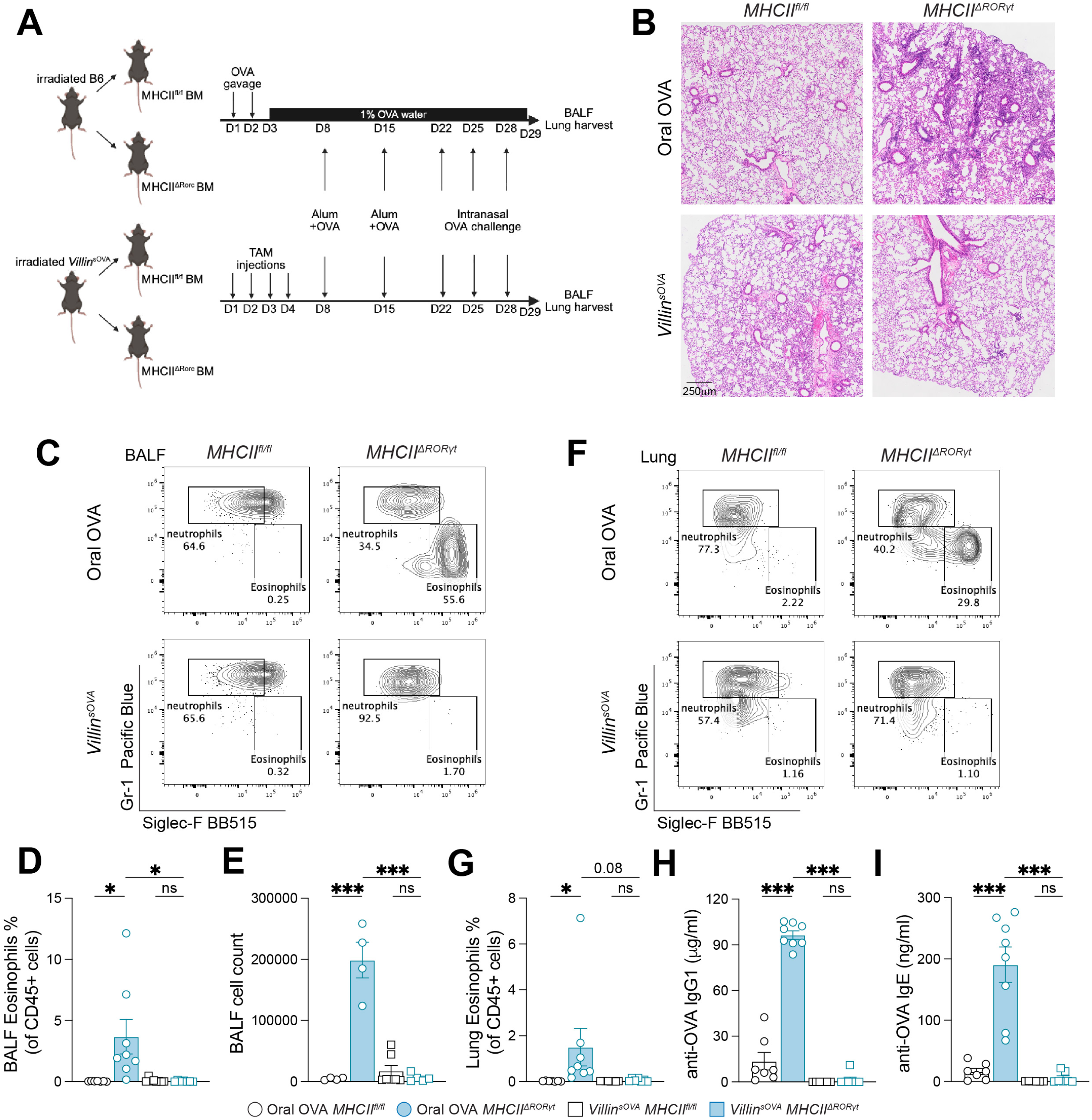
Antigen presentation by Thetis cells is dispensable for self- but not dietary antigen induced systemic tolerance. **A**. Experimental scheme for the establishment of tolerance to alum-mediated immunity. **B**. H&E staining of lung tissue from BMC receiving *MHCII^fl/fl^* or *MHCII^11RORγt^* BM fed OVA or expressing sOVA in the epithelium. The scale bar represents 250 μm. **C-D**. Flow plots of cells (**C**) and the percentage of SiglecF^+^MHCII^−^ cells (eosinophils) among total CD45^+^ cells (**D**) in BALF from mice as in B. **E-F**. Flow plots of cells (**E**) and the percentage of eosinophils among total CD45^+^ cells (**F**) in lung tissue from mice as in B. **G-H**. Serum concentration of OVA-specific IgG1 (**G**) and IgE (**H**) in mice as in B. Data are pooled from two independent experiments in D-I (*n* = 6-8 per group, as indicated by symbols). * p<0.05, ** p<0.01, *** p<0.001 by t-test.

These data show that the differential APC requirements for inducing pTreg cells specific for oral *versus* gut epithelium-derived self-antigen (Fig. 2K-N) translate into their requirement for systemic tolerance.

### Helminth infection triggers breakdown of tolerance and anaphylaxis in response to secreted but not cytosolic gut-self antigen or dietary antigen

Finally, we assessed whether the induction of Th2 cells recognizing secreted but not cytosolic self-antigen or oral antigen by *S. venezuelensis* leads to differential susceptibility to systemic allergic responses, such as elevated levels of allergic IgE and IgG1 antibodies and anaphylaxis. We exposed mice continuously to OVA either by gavage and drinking water or administration of tamoxifen to *Villin^sOVA^*or *Villin^cOVA^* mice starting from the peak of infection (day 5 post-inoculation). We compared serum anti-OVA IgE and IgG1 titers and systemic anaphylaxis (drop in body temperature and survival) following intraperitoneal OVA challenge. Uninfected mice served as controls, and the experiments were conducted both with and without OT-II transfer at the time of OVA exposure (Fig. 7A). Of all groups, only *Villin^sOVA^* mice infected with *S. venezuelensis* developed signs of anaphylaxis, manifested by a decrease in body temperature of 4°C within 30 min (Fig. 7B, Fig. S6D), markedly reduced survival (Fig. 7C), and an increase in anti-OVA IgG1 and IgE levels from almost non-detectible to levels comparable to Alum immunization^7,8^ (Fig. 7D, E). OT-II transfer produced a more severe phenotype (Fig. 7B-E), but these cells were not required, indicating that endogenous OVA-reactive CD4+ T cells were sufficient.

**Figure 7.**
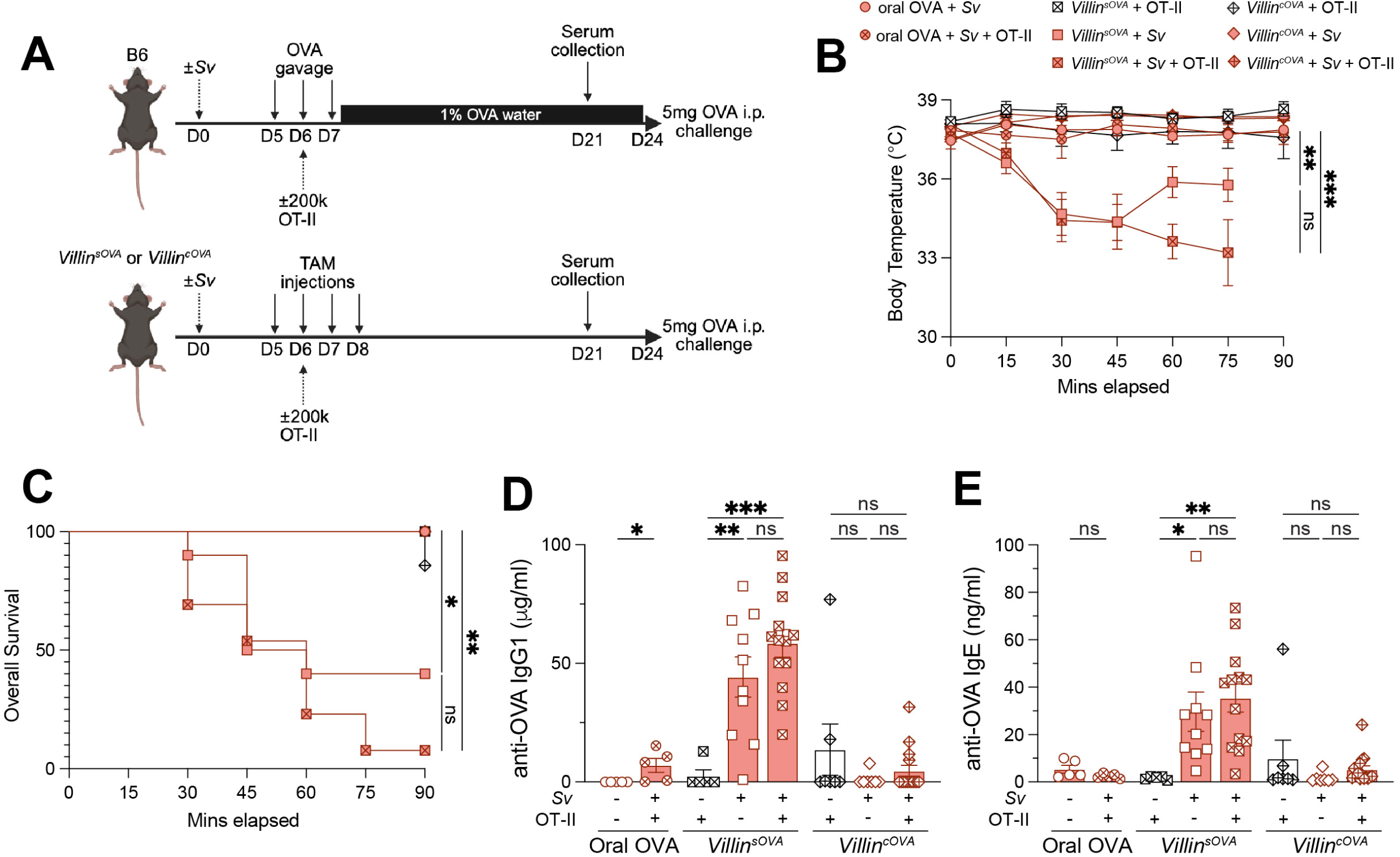
Helminth infection triggers breakdown in tolerance and anaphylaxis in response to secreted but not cytosolic gut self-antigen or dietary antigen. **A**. Experimental scheme for systemic anaphylaxis model. **B**. Body temperature measurements of indicated groups every 15 min post 5 mg i.p. OVA challenge. **C**. Anaphylaxis as measured by survival of mice at the indicated times after 5 mg i.p. OVA challenge. **D-E**. Serum concentration of OVA-specific IgG1 (**D**) and IgE (**E**) in mice as in A. Data are pooled from two independent experiments in B-E (*n* = 5-12 per group, as indicated by symbols). * p<0.05, ** p<0.01, *** p<0.001 by Logrank test and t-test.

In conclusion, these experiments demonstrate that the Th2 response to secreted, but not cytosolic or oral antigen, results in a unique susceptibility to allergic reactions.

## Discussion

In this study, we discovered that gut-derived antigens elicit different T cell fates depending on the compartment from which they originate, whether luminal (dietary and presumably bacterial) or epithelial, and if epithelial, whether secreted or cytosolic. We found that each antigen source has different APC requirements for stimulating T cell responses, which leads to corresponding disease susceptibilities (Fig. S7). Finally, by engineering Cre-inducible *OVA*-expressing mice, we developed new tools that will allow definition of the rules governing tissue-specific autoimmunity *versus* self-tolerance beyond the gut (Fig. 1A).

Our findings have major implications for understanding the general principles dictating T cell fate decisions in the gut and the potential contribution of autoimmunity to gastrointestinal diseases. The default T cell fate in the gut is Treg differentiation, with reprogramming to pro-inflammatory lineages occurring only when adjuvants or infections alter the APC profile. This mechanism likely evolved to prevent adverse reactions to food and commensal microbes while preserving host defense against pathogens or symbionts that breach the intestinal barrier, a paradigm supported by our findings. However, while responses to a single antigen like dietary OVA can be pleiotropic, and OVA-specific T cells interact with many APC subtypes when OVA is a dietary antigen^2^, we found that differential antigen access by distinct APCs is another critical and previously underappreciated determinant of T cell polarization. This became evident especially upon perturbations, where the immune system deviates from the pTreg cell default. Thus, the logic for predicting T cell fate is not simply determined by which adjuvants (“signal 3”) are present but also dictated by the dominant APC subset with access to the given antigen.

This mechanism connects to another key concept from our study: access to an antigen is determined by antigen biology- whether it is cytosolic and accessible only following cell death, secreted and readily available, and, while not formally tested in our study, potentially whether it is intact or partially degraded, or more apical or basolateral. Each type of antigen may be preferentially sampled by different APC subset repertoire, thereby shaping the ensuing T cell responses. Whether antigen access is regulated by anatomical location of the antigen relative to an APC or by cargo-selective endocytic machineries remains to be determined. While we observed this phenomenon using self-antigens, it is likely that the same rules apply to bacterial and dietary antigens that directly interact with the epithelium, such that these antigens can behave like intracellular or secreted OVA rather than like luminal antigens. For T cell responses to bacteria, which appear to be diverse even when elicited by commensals, this mechanism suggests that in addition to the adjuvant provided by a given bacterial strain, the nature of the bacterial interaction with the epithelium and consequently the APC that presents the antigen contributes to determining the dominant T cell fate in response to that strain. Finally, these concepts of APC access and antigen biology contributing to shaping the nature of the T cell response also likely apply to other tissues, particularly those at barrier sites such as the lung and skin that also regularly encounter foreign antigens.

Another striking discovery made in our experiments is that even baseline pTreg cell induction is not universally dependent on one APC subtype. Instead, the main APC required for pTreg cell induction varies with antigen source. TCs are strictly required for developing pTreg cells that recognize dietary and commensal bacterial antigens^8–10^. However, this is not the case for pTreg cells recognizing self-antigens, for which antigen presentation by TCs plays a minor role, contributing only when secretory antigen is highly abundant (Fig. 2K). This finding suggests that TCs have access to luminal antigens in a manner that does not extend to self-antigens. TCs are thought to be largely LN-resident, and therefore likely acquire antigens *in situ* rather than by migration from the gut. It is possible that their exposure to dietary and bacterial antigens, which are at least partially digested, occurs through delivery in acellular lymph, whereas secreted self-antigens may only spill into this compartment once the antigen-loading capacity of gut cDCs is saturated. Conversely, such a dual-antigen access system for pTreg cell induction may in itself support the tolerogenic default to dietary and benign commensal antigens by also enabling a temporally controlled dominance of tolerance: Postprandial lymph and lymph-borne antigens reach the LNs within 30-60 min after ingestion, while DC migration can take up to 24 h. Future research on the role of digestive enzyme-mediated antigen preprocessing, as well as antigen state and concentration in lymph, will be required to test these hypotheses. Similarly, how pTreg cell induction by TCs is suppressed in the context of infections remains unresolved. In addition to the potential reprogramming of TCs, our data suggest that antigen could alternatively be diverted to migratory DCs in such contexts (Fig. 4), but this idea also requires additional experimental confirmation. Finally, these rules may differ for pTreg cell induction outside the gut, where antigen processing by digestive enzymes is absent and lymph production and flow rates are lower.

Our data also imply that gut-self recognizing Th1/T_cyt_ cells contribute to autoimmune gut pathology more than previously appreciated. Though *bona fide* gut autoimmunity beyond CD is rare, it may in fact be underrecognized, as self-reactivity is not routinely monitored and the rapid turnover of intestinal epithelial cells may obscure its detection, particularly in the context of inflammatory bowel disease or immune checkpoint inhibitor-induced colitis, where T cell reactivity is currently thought to be directed toward the microbiota. While such self-recognizing Th1/T_cyt_ cells in the gut may be detrimental in the presence of pro-inflammatory cues such as tissue damage, these cells could provide a beneficial first line of defense against a previously unencountered type 1-inducing pathogen through the secretion of inflammatory cytokines. Similarly, secreted self-antigen-elicited Th2 cells could contribute to type 2 responses in the gut. Analogous to control of Th1/T_cyt_ cells, there likely are mechanisms to ensure that such cells do not cause damage in the absence of local gut insults. Nonetheless, it is conceivable that the persistence of these cells has been evolutionarily favored to allow rapid reinvigoration during new type 2-inducing infections. Experimentally dissecting these potentially beneficial gut self-recognizing T helper cells is another important future direction.

The findings made here challenge paradigms of how pTreg cell induction is governed in the gut and offer a unifying framework to explain how different T cell fates arise in response to intestinal antigens. Our work places self-antigens into the broader context of gut antigen recognition and establishes a universal system to study autoantigens in tissue-specific immunity.

## Limitations of the study

Most of our experiments rely on a single antigen, OVA, and cognate T cell clones, OT-I and OT-II cells. Epithelial cells are heterogeneous, and *Vil^Cre^* drives *OVA* expression in all subtypes, making it unclear which epithelial population drives the observed phenotypes. APC access to OVA from different sources is inferred indirectly through the impact of APC subtype loss or lack of presentation via MHC-II on T cell fates known to depend on each APC type. Finally, we only show that antigen presentation by TCs is not required for the generation of self-reactive pTreg cells, but do not formally rule out the possibility that TCs contribute to this process through other mechanisms.

## Acknowledgements

We thank the University of Chicago Animal Resource Center and microscopy, flow cytometry, functional genomics cores. We thank Bana Jabri and members of the Esterházy lab for invaluable discussions and suggestions along the study, and Steven Erickson for advice on engineering inducible *OVA* mice. This work was funded by the National Institutes of Health (NIH) R01 DK133393, NIH R01 AI038339, the Searle Scholar’s Program, Pew Charitable Trust, and University of Chicago start-up funds (to D. Esterházy). Additional support was provided by NIH T32 AI007090 (to M. R. Komnick), the Tull Family Foundation (G. M. Taylor), and the Heinz Endowments (to T. S. Dermody).

## Author contributions

Conceptualization, Y.Z., H.B. and D.E.; Methodology, Y.Z., H.B., E.S., M.R.K. and D.E., Investigation, Y.Z., H.B., E.S., G.T., K.F., M.R.K., S.L. and D.E.; Validation, Y.Z., H.B., E.S., M.R.K., and D.E.; Formal Analysis, Y.Z., E.S., M.R.K. and D.E.; Data Curation, Y.Z.; Writing – Original Draft, D.E.; Visualization, Y.Z.; Writing – Review & Editing, Y.Z., H.B., E.S., G.T., K.F., M.R.K., T.S.D. and D.E.; Funding Acquisition, T.S.D. and D.E.; Resources, T.S.D. and D.E; Supervision, D.E.

## Materials and Methods KEY RESOURCES TABLE

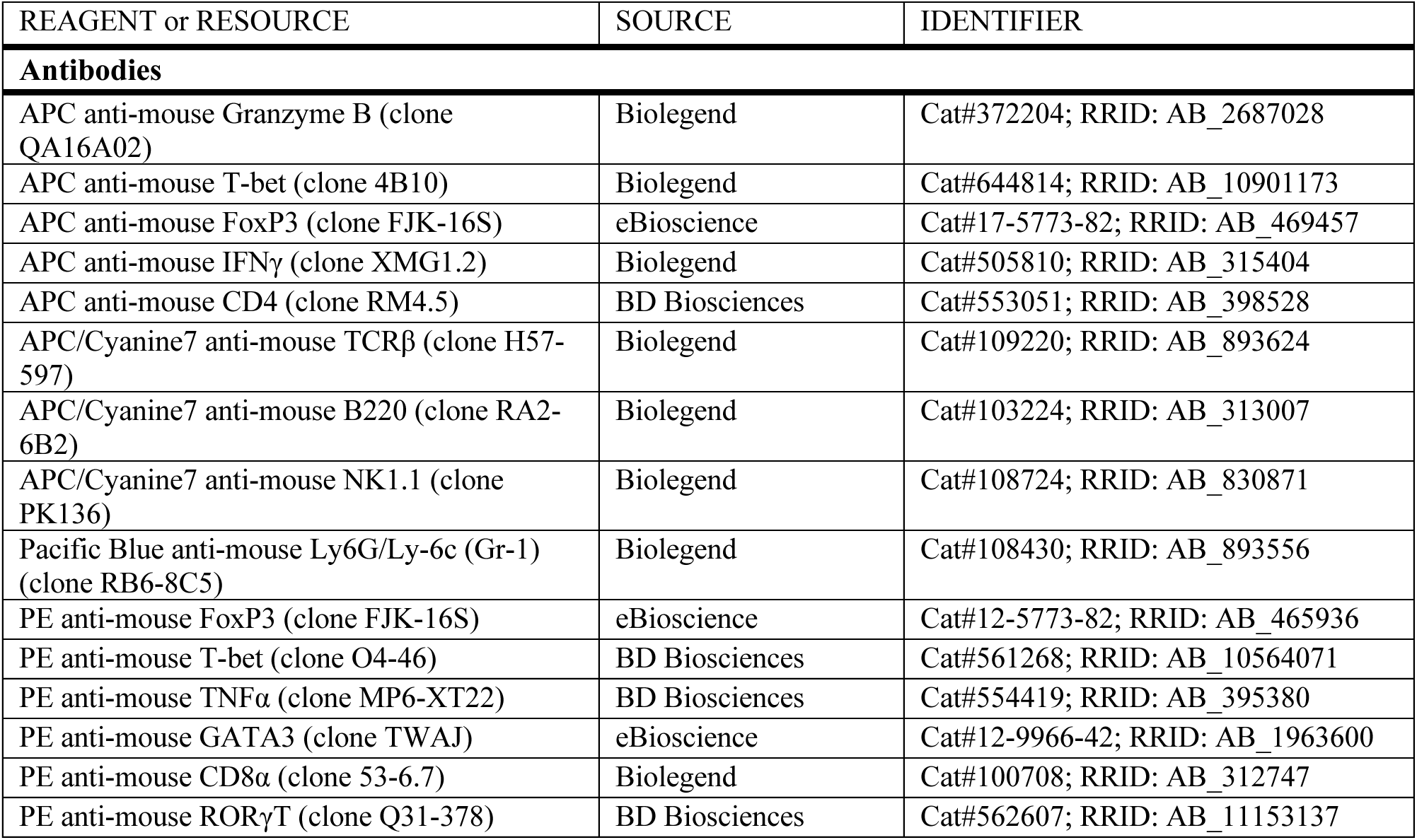
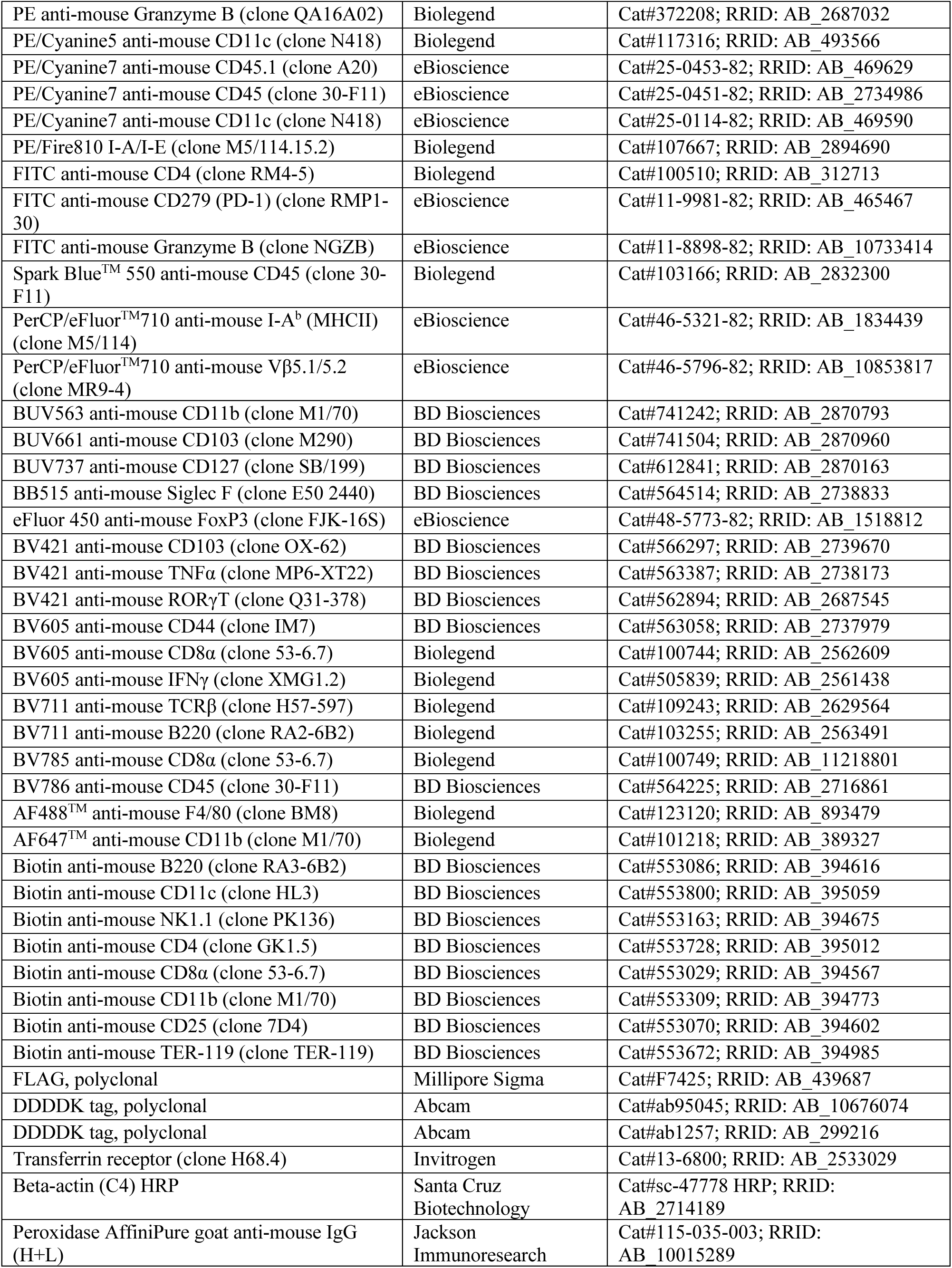
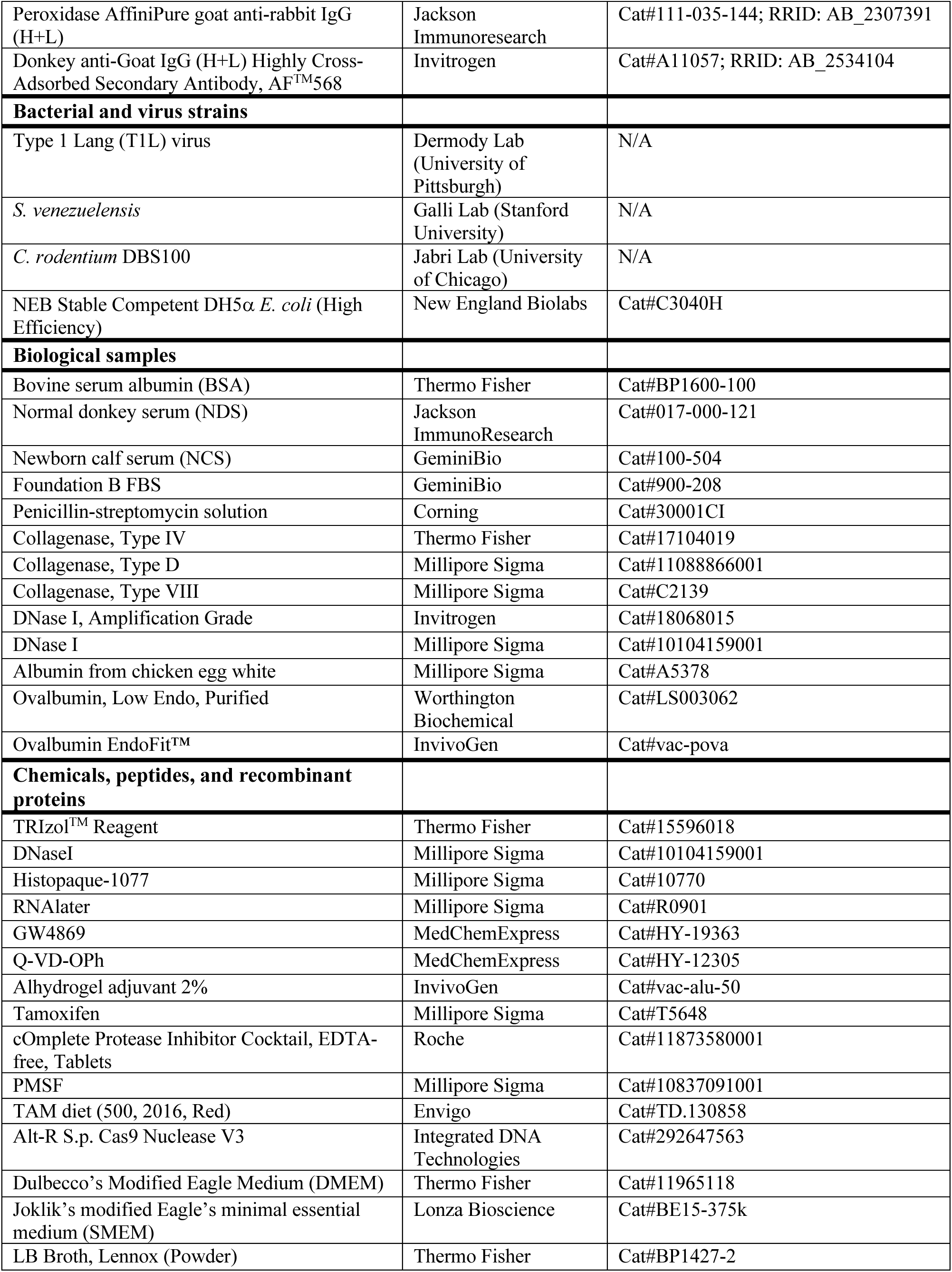
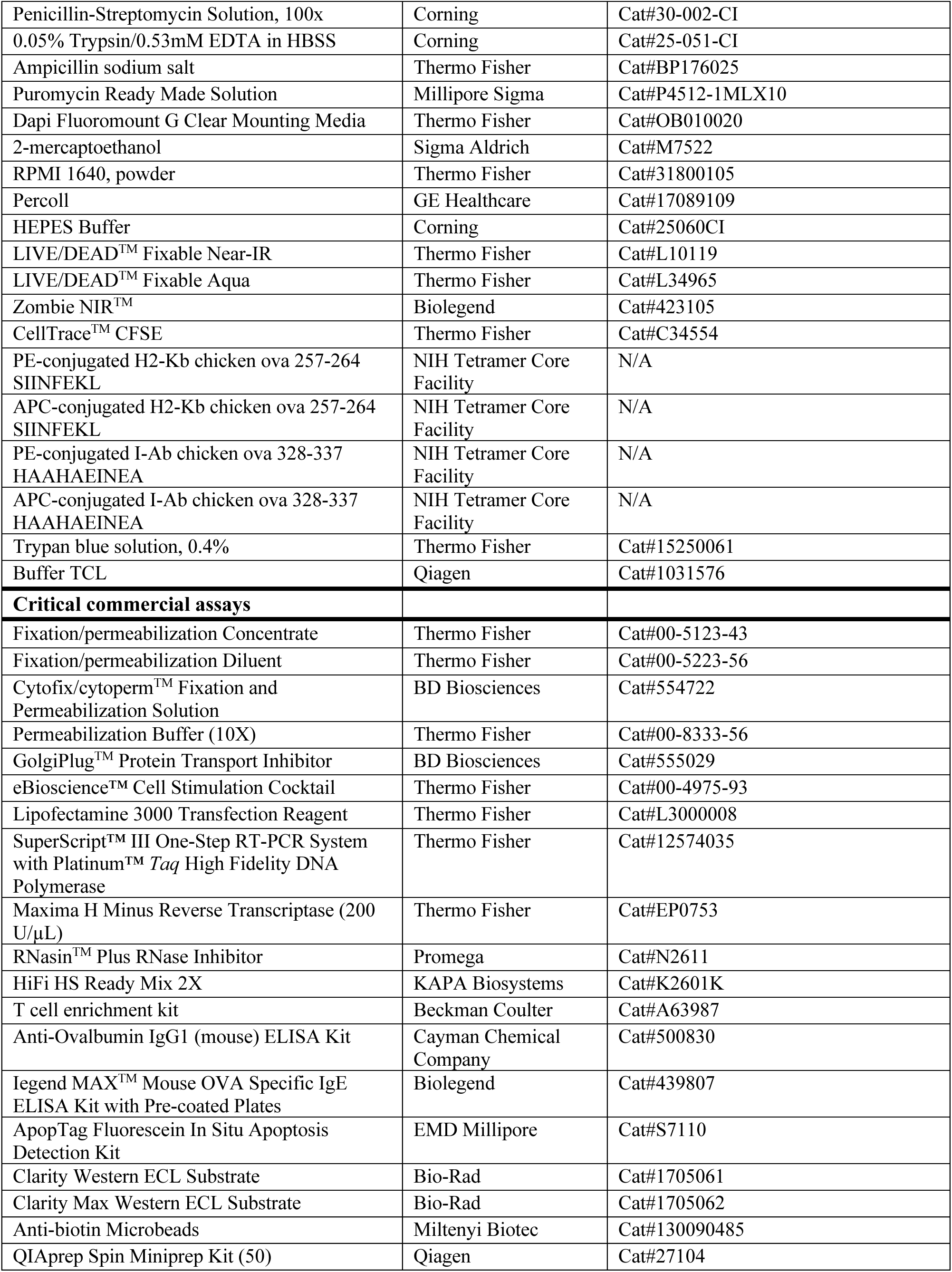
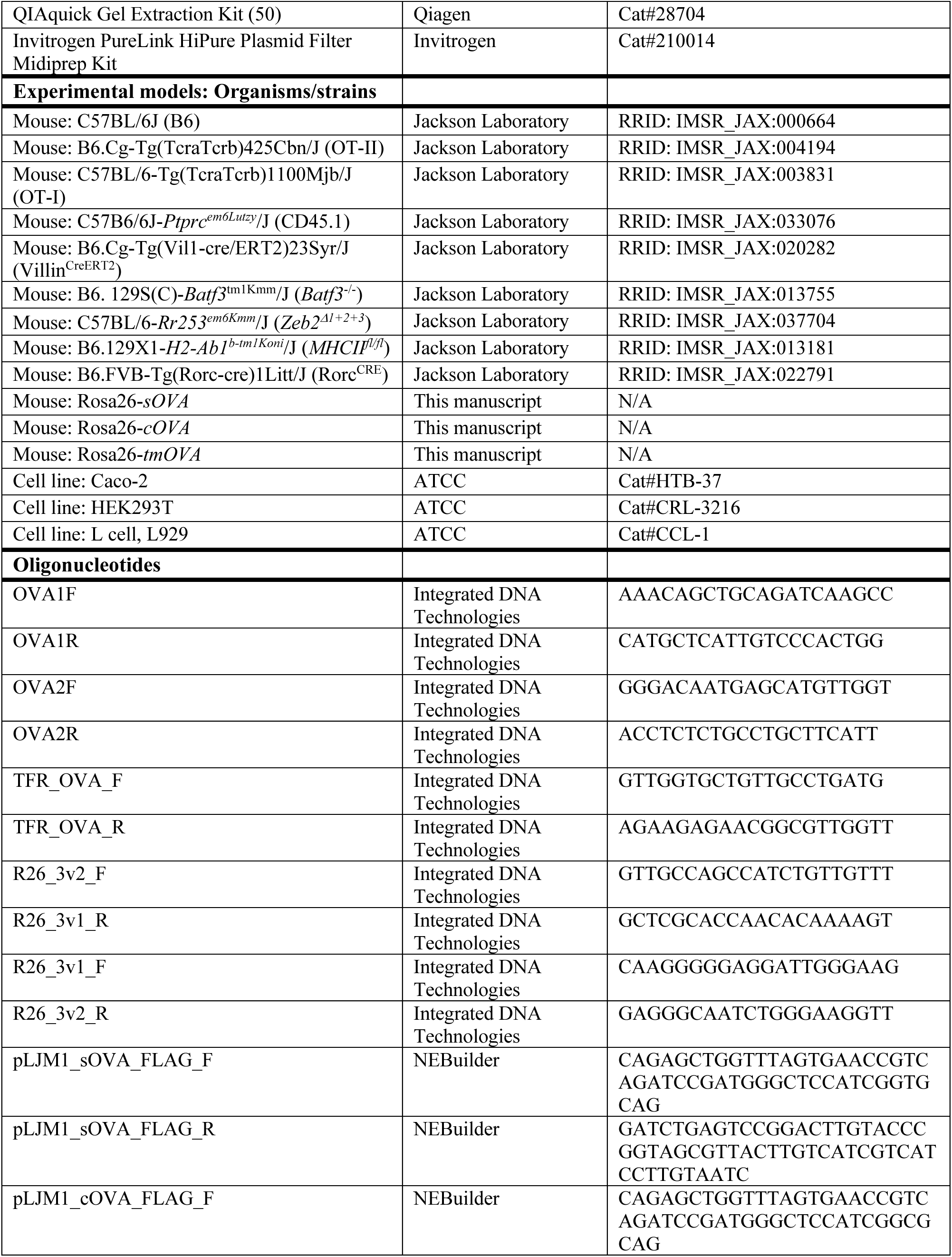
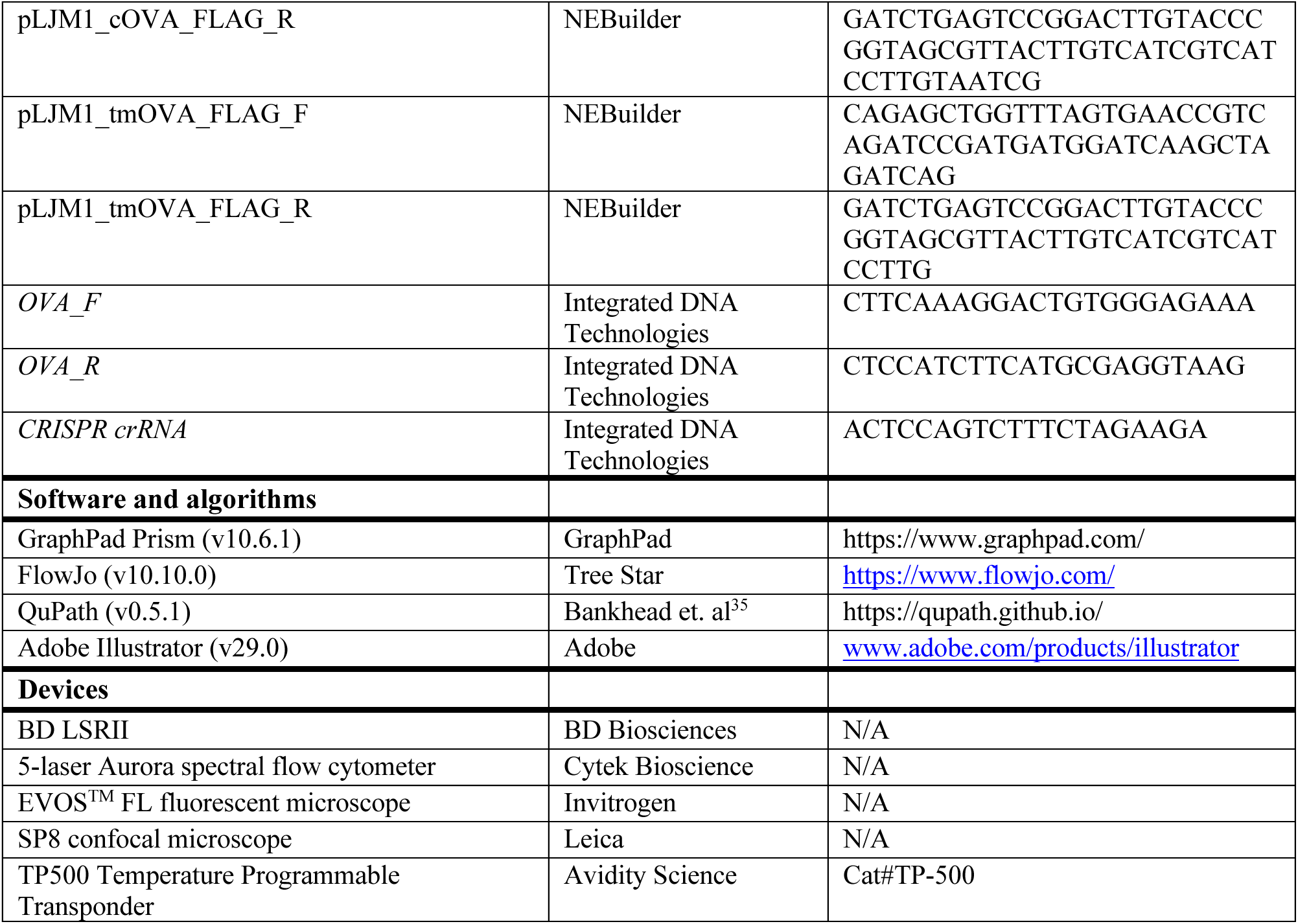

### Mice

C57BL/6J, CD45.1 congenic (C57B6/6J-*Ptprc^em6Lutzy^*/J), *Villin^CRE-ERT2^* (B6.Cg-Tg(Vil1-cre/ERT2)23Syr/J), *Batf3*^-/-^ (B6. 129S(C)-*Batf3*^tm1Kmm^/J), *Zeb2^Δ1+2+3^* (C57BL/6-*Rr253^em6Kmm^*/J), *Rorc^CRE^* (B6.FVB-Tg(Rorc-cre)1Litt/J), and *MHCII^fl/fl^*(B6.129X1-*H2-Ab1^b-tm1Koni^*/J), B6.Cg-Tg(TcraTcrb)425Cbn/J (OT-II) and C57BL/6-Tg(TcraTcrb)1100Mjb/J (OT-I) mice were purchased from The Jackson Laboratory and maintained at the University of Chicago animal facility under specific pathogen-free (SPF) conditions, confirmed to be free of segmented filamentous bacteria. OT-I and OT-II mice were crossed to CD45.1^+/+^ mice to introduce a congenic marker. All experiments were performed in accordance with AICUC protocols.

### Engineering *OVA* expression vectors and Rosa26 targeting constructs

Full length *ovalbumin* (*OVA*) and transmembrane *OVA* were amplified from Addgene plasmids #64599 and #64600, respectively, and cloned into pcDNA3.1 upstream of a C-terminal 3xFLAG tag using the In-Fusion cloning kit (Takara) with *NotI* as the restriction site. Cytosolic *OVA*, lacking amino acids 18-143 relative to the full-length sequence, was amplified from a construct provided by Seungmin Hwang and similarly inserted into pcDNA-3xFLAG. Plasmids were introduced into chemically competent *E. coli* DH5α by heat-shock transformation. Transformed bacteria were plated onto LB plates supplemented with ampicillin. Single colonies were cultured, expanded and plasmids extracted by Miniprep (Qiagen). Expression of each OVA variant was validated by transient transfection of HEK293T cells using Lipofectamine 3000 Transfection Reagents (Thermo Fisher) followed by immunoblotting.

For *Rosa26* targeting, the *OVA*-3xFLAG sequences were PCR-amplified from the respective expression construct and cloned into the *MluI/AsiSI* site of the pR26-CAG-AsiSI/MluI (Addgene #74286). All constructs were verified by Sanger sequencing. Plasmids were introduced into chemically competent *E. coli* DH5α by heat-shock transformation. Single colonies harboring pR26 targeting constructs were selected using ampicillin-supplemented LB plates and cultured overnight (≤12 h). Plasmid DNA was prepared using anion-exchange midi columns (Invitrogen). Purified DNA was ethanol-precipitated and resuspended in nuclease-free water. On the injection day, the CRISPR injection mix was prepared 2 h prior to use^18^. Briefly, 5 μl crRNA (5’-ACTCCAGTCTTTCTAGAAGA, 1μg/μl; IDT) and 10μl tracrRNA (1μg/μl; IDT) were annealed in a thermocycler (95°C with a ramp down to 25°C at 5°C/min). Cas9 protein (10 μl, 1 μg/μl; IDT) was then complexed with 10 μl of annealed crRNA:tracrRNA and 30 μl of water for 15 min at 25°C. Midi-prepped plasmid DNA was added to the RNP complex at a final concentration of 15 μg/μl in a total volume of 100 μl. The mixture was spun at 20,000 x g for 20 min, and the upper 65 μl was transferred to a fresh tube and provided to the University of Chicago Transgenic Mouse Facility.

Fertilized C57BL/6 oocytes were microinjected with the mixture and reimplanted into pseudopregnant CD-1 females. Tail biopsies were collected from pups at 2 weeks of age for genotyping. Integration of *OVA* sequences was confirmed by PCR using primers OVA1F, OVA1R, OVA2F, OVA2R, TFR_OVA_F and TFR_OVA_R. Correct targeting to the *Rosa26* locus was validated with primer pairs R26_3v2F and R26_3v1R, followed by R26_3v1F and R26_3v2R. The resultant PCR products were analyzed by agarose gel electrophoresis to confirm expected size and subsequently validated by Sanger sequencing. Successful founders were isolated in quarantine for SFB removal and bred to C57BL/6 mice for two generations before crossing onto Cre driver lines. The sequences of mentioned primers are listed in the Key Resource Table.

### Induction of ovalbumin self-antigen in the intestine by tamoxifen administration

Rosa26-*OVA* mice were crossed to *Villin^CRE-ERT2^* mice. Offspring were administrated tamoxifen-containing diet (TAM diet, Envigo) for 7 consecutive days or by daily intraperitoneal injection of 1mg tamoxifen (TAM) (Sigma) dissolved in 100 μl corn oil for 5 consecutive days at 4-6 weeks of age to induce for OVA self-antigen expression in the intestinal epithelium. Following TAM administration, mice were returned to a normal Chow diet for 4-7 days before use in experiments.

### Oral ovalbumin administration

OVA (Sigma) was diluted in sterile PBS or water, filter-sterilized and administered by oral gavage at 50mg in 200 μl PBS daily for 2-3 consecutive days. In addition, OVA was provided continuously in the drinking water at a concentration of 1% w/v until harvest.

### Segmentation of gut-draining lymph nodes

Gut-draining LNs were harvested as previously described^1^. Briefly, the LNs that are relevant for this study include: the hepatic-pancreatic (liv) LN that have little drainage from the gut as a negative control, the hepatic–coeliac (cel) LN along the portal vein, the pancreatic–duodenal (duo) LN draining the duodenum, the upper portion of the main mLN chain (jej) draining the distal duodenum and jejunum, the lower portion of the main mLN chain (ile) draining the ileum, as well as a separate LN (cec) at end of the mLN chain that drains the caecum and colon.

### Adaptive T cell transfer and CFSE tracing

Naïve T cells were isolated from pooled LNs and spleens by negative selection using biotinylated antibodies against NK1.1, B220, CD11c, CD11b, TER119 and either CD8α or CD4, followed by anti-biotin MACS beads (Miltenyi Biotec). Purity of transgenic cells was confirmed by flow cytometry (CD45.1^+^Vβ5^+^TCRβ^+^CD4^+^ for OT-II cells or CD45.1^+^Vβ5^+^TCRβ^+^CD8α^+^ for OT-I cells). Cell viability was determined by Trypan blue staining.

For experiments probing T cell outcomes in the dLNs, OT-II cells were labeled with CFSE Cell Proliferation Kit (Life Technologies) according to the manufacturer’s protocol, incubating cells for 3-5 mins at 37°C, followed by a wash in PBS. A total of 750,000 OT-II or OT-I cells was transferred by retro-orbital injections under isoflurane anesthesia, and LNs harvested 3 (after infections) or 4 (at homeostasis) days post-transfer, as indicated. For experiments probing T cell responses in the intestine, transgenic cells were not CFSE-labeled. On the day of transfer, 200,000 OT-II and 50,000 OT-I cells were injected retro-orbitally. Intestines were procured 7 days post-transfer.

### Immunofluorescence staining

Tissues were fixed in 4% PFA/PBS for 2 h at RT and subjected to paraffinization. Paraffin-embedded tissue sections (5μm) were heated at 65°C for 20min, deparaffinized twice in xylene (10 min each), and rehydrated through a graded ethanol series. Sections were permeabilized with 0.1% Triton X-100, blocked for 30 min at RT with blocking buffer (5% donkey serum, 2.5% BSA in PBS) before overnight incubation at 4°C in a humidified chamber with primary antibodies in blocking buffer (see Antibodies section). The following day, slides were washed three times with PBST (PBS with 0.05% Tween-20) for 15 min each, incubated with appropriate secondary antibodies in blocking buffer for 1 h at RT, washed again three times in PBST, and mounted with DAPI Fluoromount G Clear Mounting Media (SouthernBiotech). Images were acquired using an SP8 Lightning confocal microscope (Leica) at the University of Chicago Microscopy Core.

### Immunoblotting

Whole-cell or whole-tissue lysates were prepared in RIPA lysis buffer (20 mM HEPES, 0.2 mM EDTA, 300 mM NaCl, 1.5 mM MgCl_2_, 1% Triton X-100) supplemented with protease inhibitor (Roche) and 1 mM phenylmethylsulphonyl fluoride (PMSF; Sigma). Supernatant from cultured cells were concentrated using Amicon Ultra Centridugal Filter, 10kDa MWCO (Millipore) and mixed with Laemmli buffer. Samples were separated by SDS-PAGE and transferred onto nitrocellulose membranes (Millipore). Membranes were probed with the indicated primary and secondary antibodies (see Key Resource Table).

### Isolation of cells from lymph nodes

For T cell analysis, LNs were dissected into RPMI supplemented with 2% NCS and 1mM EDTA, disrupted using two frosted glass slides, and filtered into 96-well plates. Cells were immediately subjected to Live/Dead and surface marker staining. For APC analysis, LNs were dissected into RPMI supplemented with 2% NCS and 1% HEPES, finely minced, and digested with 2.5 mg/ml Collagenase D (Roche) for 30 min at 37°C. Resulting single-cell suspensions were directly used for Live/Dead and surface marker staining.

### Isolation of lymphocytes from small and large intestines

The small intestine was excised, cleared of mesentery, Peyer’s patches and fecal contents. Intestines were cut longitudinally and washed three times in PBS containing 1μM DTT to remove mucus. Tissue was cut into ∼1cm pieces and incubated in PBS with 30 mM EDTA for 10 min at 37°C with shaking at 230 rpm, followed by a vigorous shake after the incubation. Tissue samples were filtered through a metal sieve and the flow through was collected as intraepithelial fraction (IEL). Tissues were transferred to a new tube containing fresh PBS without EDTA and incubated for another 10 min at 37°C at 230 rpm. Tissues were washed with PBS on a metal sieve, finely chopped and digested in RPMI containing 2% FCS, 1% HEPES, and 2 mg/ml Collagenase 8 (Roche), 200 μg/ml DNase I (Roche) for 30 min at 37°C with gentle agitation at 80 rpm. Digests were quenched in cold RPMI containing 2% FCS and cell pellets were resuspended in 40% Percoll. Lymphocytes were enriched at the interphase of a discontinuous Percoll gradient (80%/40%) by centrifugation at 2300 rpm for 23 min at RT with slow acceleration and deceleration. Cells were washed and used for flow cytometry analysis.

### T cell stimulation

For the detection of OT-I cytokine production, single cell suspensions of isolated lymphocytes were pelleted by centrifugation and resuspended in T cell media containing cell stimulation cocktail (1:500; eBioscience). Cultures were incubated for 4 h at 37°C. Following stimulation, cells were pelleted, washed once with fresh medium, and processed for cell staining.

### Flow cytometry

Single-cell suspensions were first stained with LIVE/DEAD™ Fixable Near-IR Dead Cells Stain kits (Invitrogen) in PBS for 10 min at 4°C, followed by surface staining with antibody cocktails (in 1:200 dilution) in FACS buffer (PBS, 1% BSA, 2 mM EDTA) for 20 min at 4°C. For intracellular cytokine detection, cells were fixed and permeabilized with Cytofix/cytoperm™ solution (BD Biosciences) for 20 min prior to staining with intracellular antibodies overnight at 4°C. For transcription factor analysis, cells were incubated in Fixation/permeabilization solution (eBiosciences) for 30 min to 2 h, and subsequently stained with nuclear antibodies overnight at 4°C. All intracellular and nuclear antibodies were diluted 1:100 in 1x Permeabilization buffer (eBiosciences). Fluorescent signals were acquired on an LSR II (BD Biosciences) or a 5-laser Aurora spectral flow cytometer (Cytek) and analyzed using FlowJo software. Cell division index was calculated using FlowJo formula as described (http://www.flowjo.com/v765/en/proliferation.html), whereby the index represents the fraction of total cell divisions normalized to the estimated starting cell number.

### OVA tetramer staining

Single-cell suspensions were enriched for T cells using EasySep^TM^ Mouse T Cell Isolation Kit (Stemcell Technologies) and then incubated with Fc block for 10 min at 4°C to prevent nonspecific binding. For detection of OVA-specific CD8^+^ T cells, H-2Kb:OVA (SIINFEKL) PE- and APC-conjugated tetramers (NIH Tetramer Core Facility) were added at 1:200 in FACS buffer and incubated for 20 min at RT in the dark. For detection of OVA-specific CD4^+^ T cells, I-Ab:OVA (HAAHAEINEA) PE- or APC-conjugated tetramers (NIH Tetramer Core Facility) were added at 1:100 in T cell medium and incubated for 2 h at RT in the dark^8,36^. Cells were then washed with FACS buffer before staining with surface antibody cocktail. Each tetramer lot was validated using OVA-immunized mice before use. Tetramer-positive gates were defined using OVA-naïve mice and fluorescence-minus-one controls.

### Cell lines

HEK293T cells and Caco-2 cells were cultured in high glucose Dulbecco’s Modified Eagle Media (DMEM; Gibco) supplemented with 10% FBS and 100 U/mL penicillin/streptomycin. Spinner-adapted murine L929 cells were maintained in either suspension or monolayer cultures in Joklik’s modified Eagle’s minimal essential medium (SMEM; Lonza) supplemented to contain 5% fetal bovine serum (FBS; Gibco), 2 mM L-glutamine, 100 U/ml of penicillin, 100 μg/ml of streptomycin (Gibco), and 25 ng/ml of amphotericin B (Sigma). Cultures were maintained at 37°C in a humidified incubator with 5% CO_2_.

### Lentiviral transduction of ovalbumin expression

pLJM1-Empty (Addgene #91980) plasmid was used as lentiviral vector, which was linearized by Nhel-HF (NEB) at 37°C for 20min and gel purified. Primers were designed using NEBuilder (https://nebuilder.neb.com/#!/) to create the necessary overlap with the plasmid. Insert fragments were created using the designed primers (listed in Key Resource Table) and Phusion High-fidelity DNA polymerase (NEB). The fragments were gel purified. Gibson assembly reaction was set up and the molar ratio for insert-to-vector set to be 1:1. The reaction was incubated at 50°C for 2 h and the product was transformed into *E. coli* DH5α according to the manufacturers’ protocol. Transformed *E. coli* cells were plated onto selection agar plates and single colonies were picked and sequences. The correct colonies were expanded, and plasmids were extracted using QIAGEN Plasmid Mini Kit.

Lentivirus was produced in HEK293T cells transfected at 80-90% confluency using Lipofectamine 3000 (Invitrogen) as recommended by the manufacturer using psPAX2 (Addgene) and pMD2.G (Addgene) packaging vectors. Medium was changed 8-12 hours after transfection and supernatant was collected after 48-72 h. Viral media was passed through a 0.45μm filter and mixed with 10 μg ml^-1^ Polybrene (Sigma Aldrich) before being added to recipient cells. Infected cells were treated with puromycin to generate stable cell populations. OVA expression was verified by immunoblotting.

### Quantitative real-time PCR

Total RNA from samples was extracted using TRIzol (Thermo Fisher Scientific), treated with DNase I (Invitrogen), and reverse transcribed into cDNA using SuperScript IV kit (Thermo Fisher Scientific) according to the manufacturer’s protocol. Quantitative real-time PCR was performed using Power SYBR Green PCR Master Mix (Applied Biosystems) on Quanstudio 6 Flex machine (Thermo Fisher Scientific). The relative expression of target genes was determined by normalization to housekeeping gene *36B4* and calculated using the formula of 2^−ΔCt^ ×10000. All data were averaged from at least 2 replicates.

### Infections

Reovirus strain type 1 Lang was recovered using plasmid-based reverse genetics^37^ and purified using CsCl_2_ gradient centrifugation^38^. Titers of purified virus stocks were determined by plaque assay using L929 cells. Mice were inoculated with 10^9^ PFU of purified T1L diluted in 100 μL PBS by oral gavage^38^. OT-II or OT-I cells were transferred at the time of infection.

*S. venezuelensis* was maintained in NSG mice by subcutaneous infection with 10,000-30,000 larvae. Fecal pellets from infected NSG mice were collected and spread on Whatman paper, and incubated in water at 28°C. Larvae emerging over 3-4 days were collected to reinitiate the infection cycle or to be used for infection of experimental mice. For experimental mouse infection, 1000 larvae were administrated subcutaneously to male mice. Successful infection was confirmed by appearance of eggs in the fecal content of the infected mice. OT-II cells were transferred 5 days post infection.

*C. rodentium* was first cultured in 5 ml of LB overnight and then sub-cultured in 200 ml of LB on the next day for 4 h until OD600 reached 0.75 (exponential growth phase). The bacteria were spun down and resuspended in PBS. Mice were gavaged with 1 × 10⁹ CFU in 200 μl, and OT-II cells transferred 8 days later.

### Bone marrow chimeras

Recipient mice were lethally irradiated with two doses of 550 rads, 4 h apart, and reconstituted with 2 × 10^6^ to 5 × 10^6^ bone marrow cells delivered by retro-orbital injection under isoflurane anesthesia. Chimeras were used for experiments ≥8 weeks post-transfer.

### ApopTag staining and apoptotic index

Apoptosis was detected using ApopTag Fluorescein In Situ Apoptosis Detection Kit (Sigma). Paraffin sections were deparaffinized and rehydrated as described for immunofluorescence staining, followed by ApopTag staining according to the manufacturer’s protocol. The stained slides were scanned and analyzed in QuPath. Apoptotic index represents the percentage of ApopTag^+^ cells over total cell number.

### Q-DV-OPh *in vivo* treatment

Mice were treated with the pan-caspase inhibitor Q-VD-OPh dissolved in DMSO and diluted in sterile corn oil. A final concentration of 10 mg/kg was administered by intraperitoneal injection once daily for 5 days, starting 3 days prior to OT-II cell transfer. Control animals received vehicle (DMSO in corn oil) by the same route and schedule.

### GW4869 *in vivo* treatment

Mice were treated with exosomal biogenesis inhibitor GW4869 reconstituted in DMSO and then diluted in 300 μl sterile PBS before treatment. GW4869 was administered intraperitoneally at a dose of 2.5 mg/kg every day for the duration of the experiment, beginning 3 days prior to OT-II cell transfer. Control animals received vehicle injections on the same schedule.

### Indomethacin-induced gut injury post intestinal viral infection

Indomethacin was dissolved in sterile sodium carboxymethyl cellulose and administered at 9 mg/kg by oral gavage in a volume of 200 μl 7 days post transgenic T cell transfer and T1L infection. Mice were monitored daily for weight loss and clinical adverse events (which did not occur). Five days after treatment, small intestines were harvested, rolled into “Swiss-rolls”, fixed and Hematoxylin and Eosin (H&E)-stained for histological assessment of mucosal injury.

### Alum immunization and airway challenge

Seven days after the first oral OVA administration or first tamoxifen treatment, mice were intraperitoneally immunized with 10μg endotoxin-free OVA adsorbed to 100 μl Alhydrogel adjuvant 2% (InvivoGen) in a final volume of 400 μl PBS. Immunization was repeated after 7 days. To induce airway inflammation, mice were anesthetized and intranasally administered 10 μg endotoxin-free OVA in 50 μl PBS (25 μl per nostril) on days 14, 17, and 21 following the first intraperitoneal immunization. Mice were sacrificed one day after the last intranasal challenge.

### Bronchoalveolar lavage, lung histology, and infiltrate analysis by flow cytometry

Mice were euthanized, the trachea cannulated, and lungs were lavaged first with 0.5 ml and then 1.0 ml PBS. Total BAL cells were counted after erythrocyte lysis, and stained for flow cytometry.

Lungs were perfused via the left ventricle with 10 ml saline to remove residual blood. One lobe was digested in 400 U/ml collagenase D in HBSS for flow cytometry analysis; eosinophils were identified as CD45^+^SSC^hi^MHCII^−^CD11b^+^Ly6G^int^SiglecF^+^. Another lobe was fixed in 4% PFA, embedded in paraffin, sectioned at 5μm, and stained with H&E.

### Anti-OVA IgG1 and IgE ELISA

Mouse sera were collected and stored at -80°C until use. ELISAs were performed according to the manufacturer’s protocols using Anti-Ovalbumin IgG1 (mouse) ELISA Kit (Cayman Chemical) and Legend MAX™ Mouse OVA Specific IgE ELISA Kit (BioLegend). Samples were diluted 1:1000 for IgG1 detection and 1:3 for IgE detection.

### Allergy and systemic anaphylaxis model

Mice received 1000 *S. venezuelensis* larvae 5 days prior to the first OVA gavage or tamoxifen injection. 200,000 OT-II cells were optionally transferred at 6 days post infection. Sera were collected 21 days post infection. Mice were implanted with subcutaneous electronic temperature probes (Avidity TP-500) one or two days prior to OVA challenge. 24 days post infection, mice were then challenged intraperitoneally with 50 μg or 5 mg endotoxin-low OVA, and body temperature was recorded every 15 min for 90 min to assess anaphylactic responses. Death by anaphylactic shock was also recorded.

## Statistical analysis

All statistical analyses were performed using GraphPad Prism 10. Error bars represent the standard error of the mean (s.e.m.). For comparisons involving more than two groups, one-way ANOVA with Tukey’s multiple-comparison *post hoc* test was applied. Comparisons between two groups were performed using a two-tailed unpaired Student’s t-test assuming a Gaussian distribution. Survival curves were analyzed using log-rank tests. P-values or representative symbols are noted when differences are less than or equal to 0.1; all other differences were not found to be significant (n.s., p > 0.1). Throughout, ∗ designates p < 0.05, ∗∗ p < 0.01, and ∗∗∗ p < 0.001.

---

**Figure S1.**
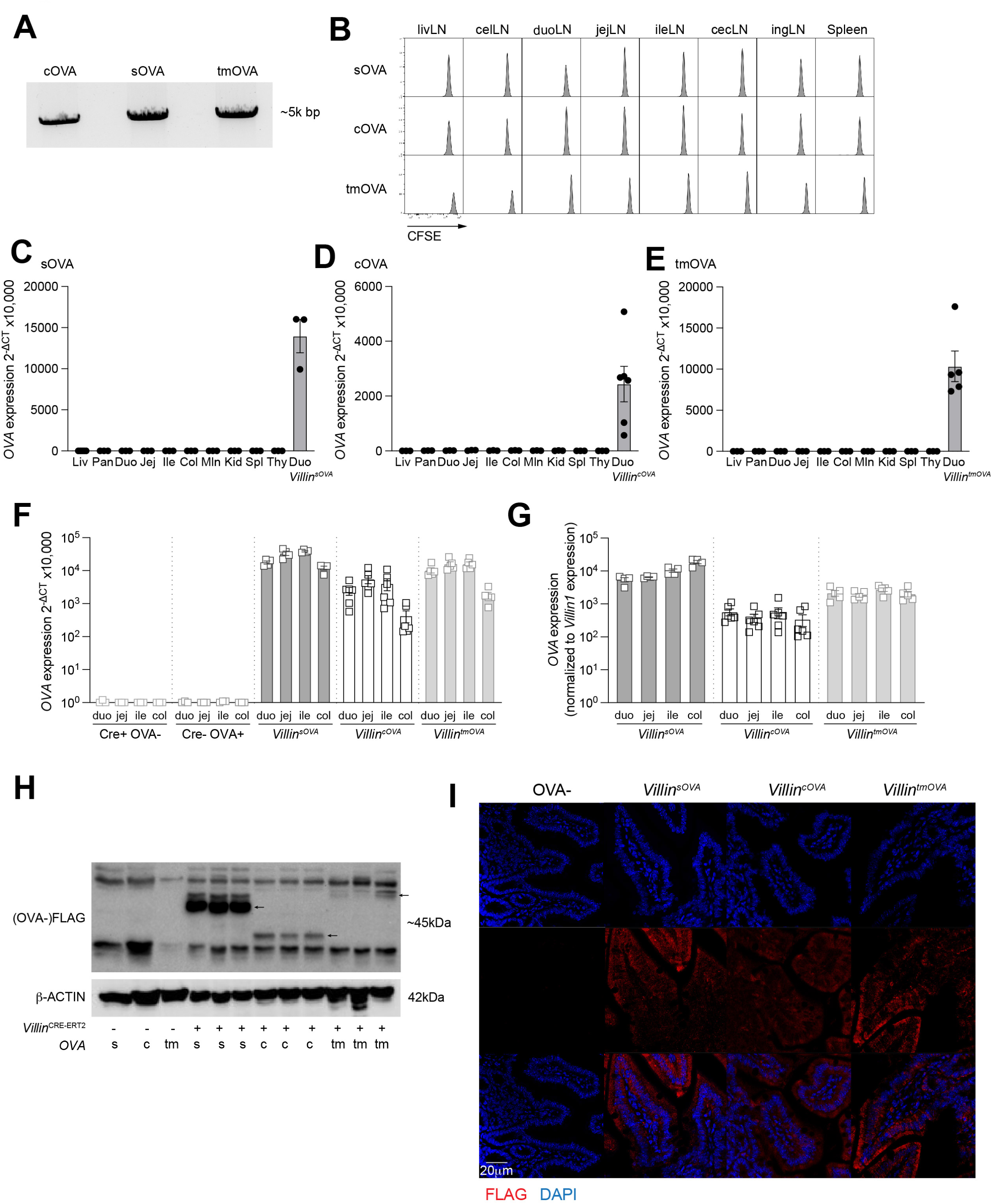
Generation and characterization of mice expressing OVA in different subcellular epithelial cell compartments. **A.** PCR amplification of the *Rosa26* locus between the homology arms from founders of each OVA line. The bands were sent for sequencing to confirm the correct sequences. **B**. CFSE dilution flow plots of OT-II cells in the indicated LNs 96 h after transfer into *sOVA^+/w^*, *cOVA^+/w^* and *tmOVA^+/w^*mice. **C-E**. *OVA* expression levels in the indicated tissues of *sOVA^+/w^*(**C**), *cOVA^+/w^* (**D**) and *tmOVA^+/w^* (**E**) mice. Duodenum from each line crossed to *Villin*^CRE-ERT2^ mice was used as a positive control (*n* = 3-6 per group, as indicated by symbols). **F-G**. *OVA* expression (**F**), and their normalization to *Villin1* (**G**), in the different gut segments of mice with indicated genotypes (*n* = 3-6 per group, as indicated by symbols). **H**. Western blot of tissue lysates of jejunum from *Villin^sOVA^*, *Villin^cOVA^* and *Villin^tmOVA^*mice and their CRE-negative controls, immunoblotted using antibodies against FLAG and β-ACTIN. Arrows indicate OVA signals. **I**. Jejunal sections from *Villin^sOVA^*, *Villin^cOVA^* and *Villin^tmOVA^* mice and their CRE-negative control stained for FLAG and DAPI (nuclei). The scale bar represents 20 μm. Data represent one experiment in C-G.

**Figure S2.**
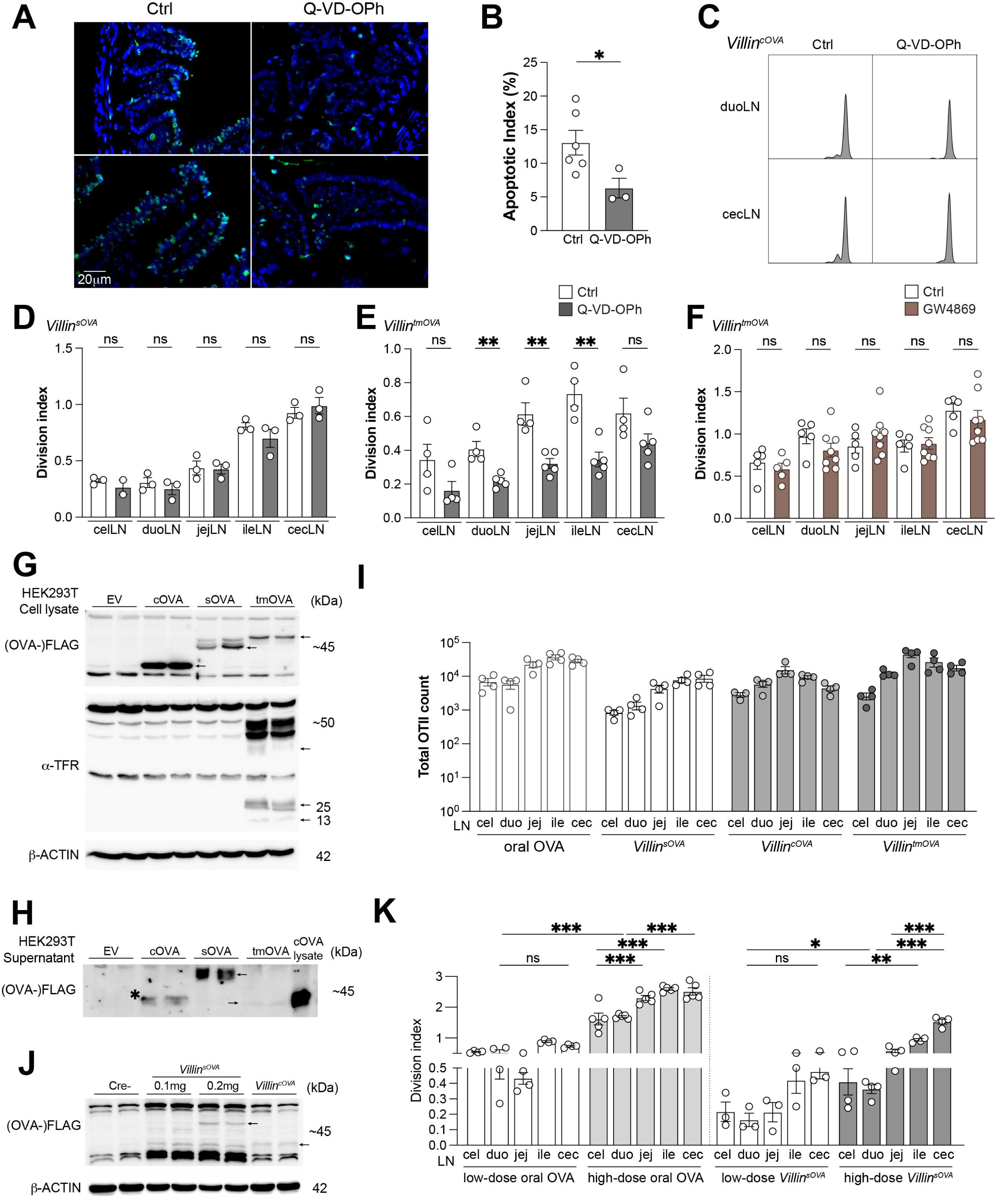
Antigens with different subcellular compartments are taken up by APCs through distinct mechanisms. **A-B.** Jejunal sections from C57Bl/6 mice treated with Q-VD-OPh or vehicle control stained for apoptotic cells and DAPI (nuclei) (**A**). Apoptotic index of A (**B**) (*n* = 3-6 per group, as indicated by symbols). **C**. Representative CFSE dilution flow plots of OT-II cells in duoLNs and cecLNs 96 h after transfer into *Villin^cOVA^* mice treated with Q-VD-OPh or vehicle control. **D-F**. Division index of OT-II cells 96 h after transfer into *Villin^sOVA^* (**D**) and *Villin^tmOVA^* (**E**) mice treated with Q-VD-OPh or vehicle control and *Villin^tmOVA^* mice treated with exosomal biogenesis inhibitor GW4869 (**F**) (*n* = 3-8 per group, as indicated by symbols). **G-H**. Western blot of cell lysates (**G**) and supernatants (**H**) of HEK293T cells transduced with lentivirus encoding empty vector, *sOVA*, *cOVA*, or *tmOVA* immunoblotted using antibodies against FLAG, α-TFR, and β-ACTIN. Arrows indicate OVA signals, asterisk possible signal from dead cells. **I.** Total OT-II cells in the indicated LNs from mice fed OVA or *Villin^sOVA^*, *Villin^cOVA^* and *Villin^tmOVA^* mice (*n* = 4 per group). **J**. Western blot of jejunal tissue lysates from *Villin^sOVA^* mice treated with low-dose of tamoxifen immunoblotted using antibodies against FLAG and β-ACTIN. Jejunal lysates from *Villin^cOVA^* mice receiving full tamoxifen diet regime were used as a reference. Arrows indicate OVA signals. **K**. Division index of OT-II cells in the indicated LNs 96 h after transfer into mice fed high- or low-dose oral OVA or mice with high or low expression of sOVA (*n* = 3-5 per group, as indicated by symbols). Data are representative of two independent experiments in B, D-F. Data represent one experiment in I, K.* p<0.05, ** p<0.01, *** p<0.001 by one-way ANOVA test and t-test.

**Figure S3.**
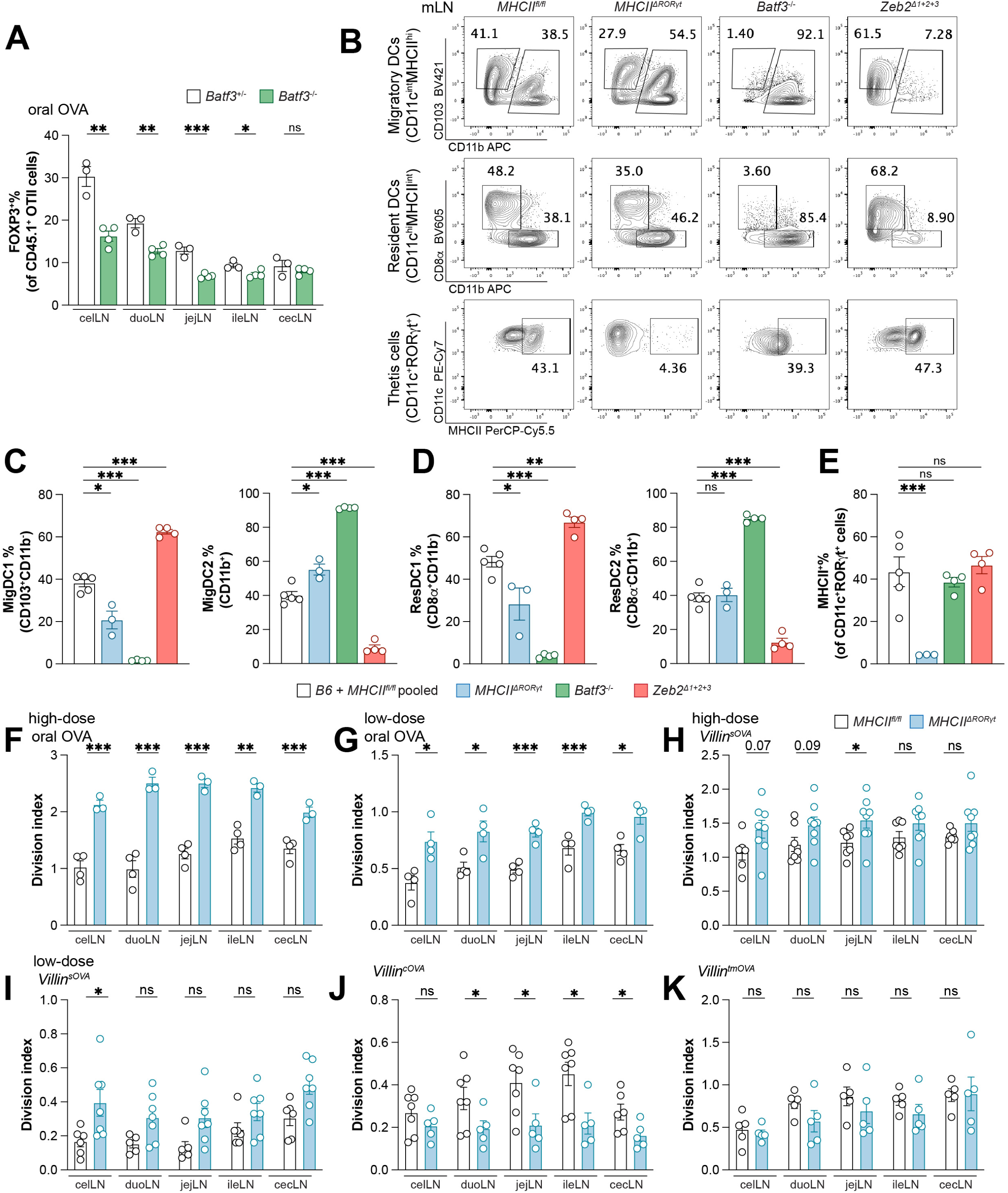
cDC1s are required while Thetis cells are dispensable for the generation of gut self-antigen specific FOXP3+ T cells. **A.** Frequencies of FOXP3^+^ among total OT-II cells in the indicated LNs 96 h after transfer into mice fed oral OVA that are *Batf3*-proficient or -deficient (*n* = 3-4 per group, as indicated by symbols). **B-E**. Representative flow plots of pooled mLNs from mice with indicated genotypes showing APC frequencies (**B**), along with quantification of migDCs (**C**), resDCs (**D**) and MHCII expression on TCs (**E**) (*n* = 3-5 per group, as indicated by symbols). **F-K**. Division index of OT-II cells in the indicated LNs 96 h after transfer into BMC receiving *MHCII^fl/fl^* or *MHCII^11RORγt^* BM fed high-dose (**F**) or low-dose dietary OVA (**G**), or mice expressing high-dose sOVA (**H**), low-dose sOVA (**I**), cOVA (**J**) and tmOVA (**K**) in the epithelium (*n* = 3-7 per group, as indicated by symbols). Data represent one experiment in A-K.* p<0.05, ** p<0.01, *** p<0.001 by t-test.

**Figure S4.**
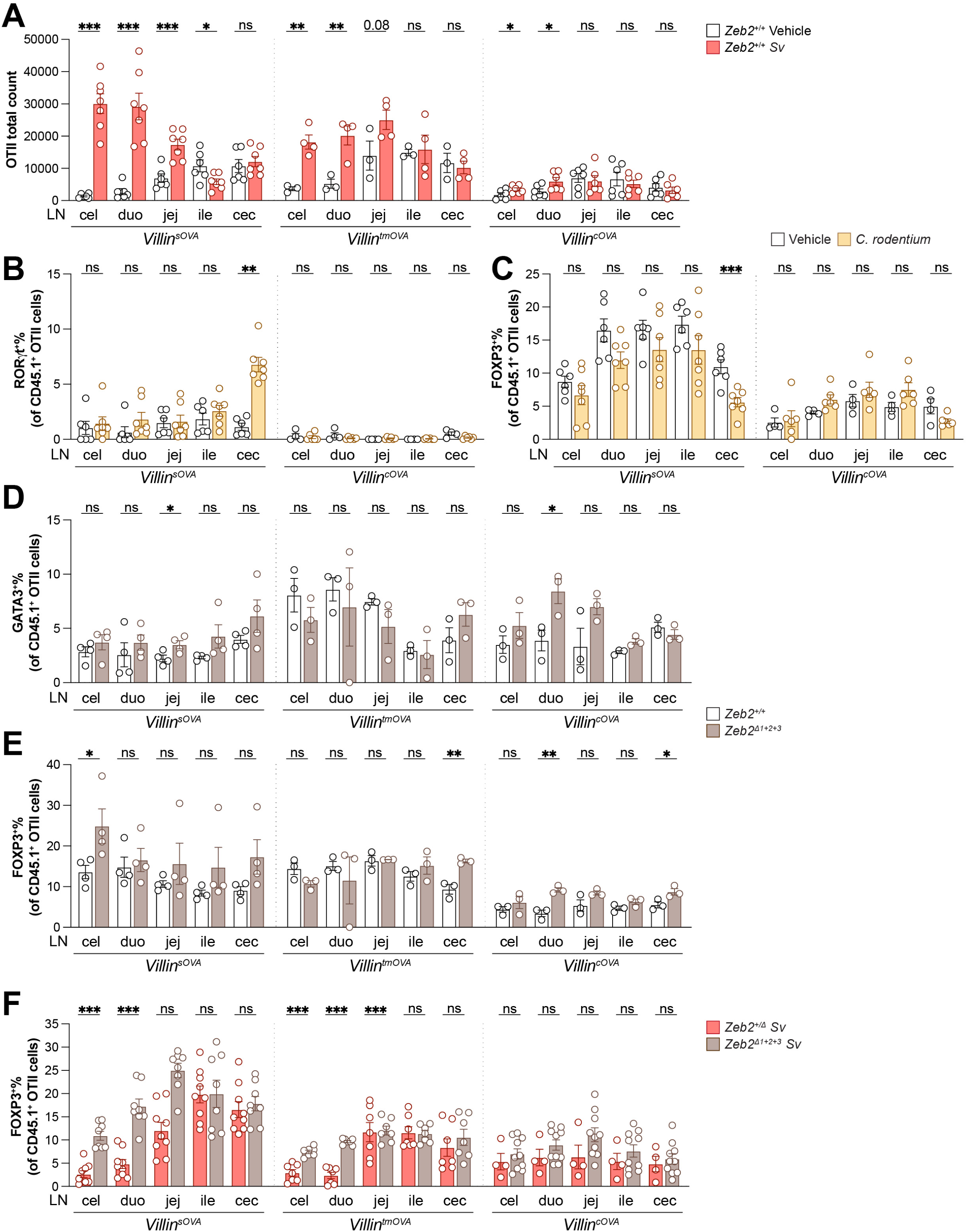
Secreted but not cytosolic self-antigens induce Th2 cells in response to helminth infection in a cDC2-dependent manner. **A.** Total OT-II cells in the indicated LNs 72 h after transfer into *Villin^sOVA^*, *Villin^tmOVA^* and *Villin^cOVA^*mice that are *S. venezuelensis*- or vehicle-infected (*n* = 3-7 per group, as indicated by symbols). **B-C.** Frequencies of RORγt^+^ (**B**) and FOXP3^+^ (**C**) among total OT-II cells in the indicated LNs 72 h after transfer into *C. rodentium*-infected *Villin^sOVA^* and *Villin^cOVA^* mice (*n* = 4-7 per group, as indicated by symbols). **D-E.** Frequencies of GATA3^+^ (**D**) and FOXP3^+^ (**E**) among total OT-II cells in the indicated LNs from bone marrow chimera (received *Zeb2^+/+^* or *Zeb2^111+2+3^* bone marrow) mice expressing OVA (*n* = 3-4 per group, as indicated by symbols). **F.** Frequencies of FOXP3^+^ among total OT-II cells 72 h after transfer into *S. venezuelensis*-infected *Villin^sOVA^*, *Villin^tmOVA^* and *Villin^cOVA^* mice proficient or deficient for *Zeb2* enhancer (*n* = 4-10 per group, as indicated by symbols). Data are representative of two independent experiments in A-E. Data are pooled from two independent experiments in F. * p<0.05, ** p<0.01, *** p<0.001 by t-test.

**Figure S5.**
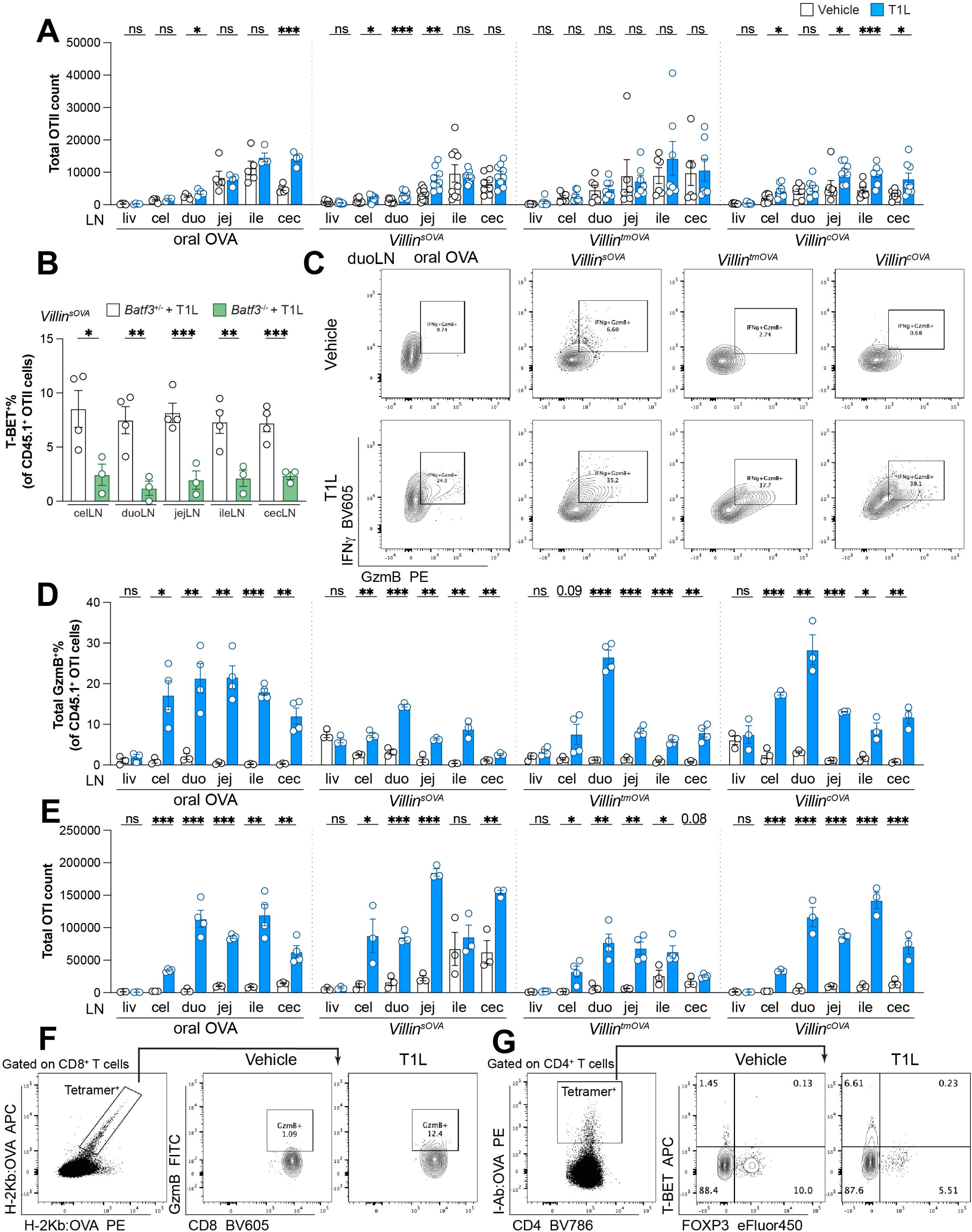
Intestinal viral infection impacts epithelial self-antigens and oral antigen-specific T cell fate similarly. **A.** Total OT-II cell number in indicated LNs of T1L- or vehicle-infected mice that were fed OVA or express sOVA, tmOVA or cOVA in the epithelium (*n* = 4-9 per group, as indicated by symbols). **B.** Frequencies of T-BET^+^ OT-II cells in the indicated LNs 72 h after transfer into T1L- or vehicle-infected *Villin^sOVA^* mice that were *Batf3*-proficient or -deficient (*n* = 3-4 per group, as indicated by symbols). **C.** Flow plots of OT-I cells in duodenal LNs of T1L- or vehicle-infected mice that were fed oral OVA or express sOVA, tmOVA or cOVA in the epithelium. **D-E.** Frequencies of GzmB^+^ among total OT-I cells (**D**) and total OT-I cell number (**E**) in the indicated LNs 72 h after transfer into T1L- or vehicle-infected mice that were fed OVA or express sOVA, tmOVA or cOVA in the epithelium (*n* = 3-4 per group, as indicated by symbols). **F-G.** Flow plots of H-2Kb:OVA^+^ (**F**) and I-Ab:OVA^+^ (**G**) cells in total mLNs of T1L- or vehicle-infected *Villin^sOVA^* mice. Data are pooled from two independent experiments in A. Data represent one experiment in B, D, E. * p<0.05, ** p<0.01, *** p<0.001 by t-test.

**Figure S6.**
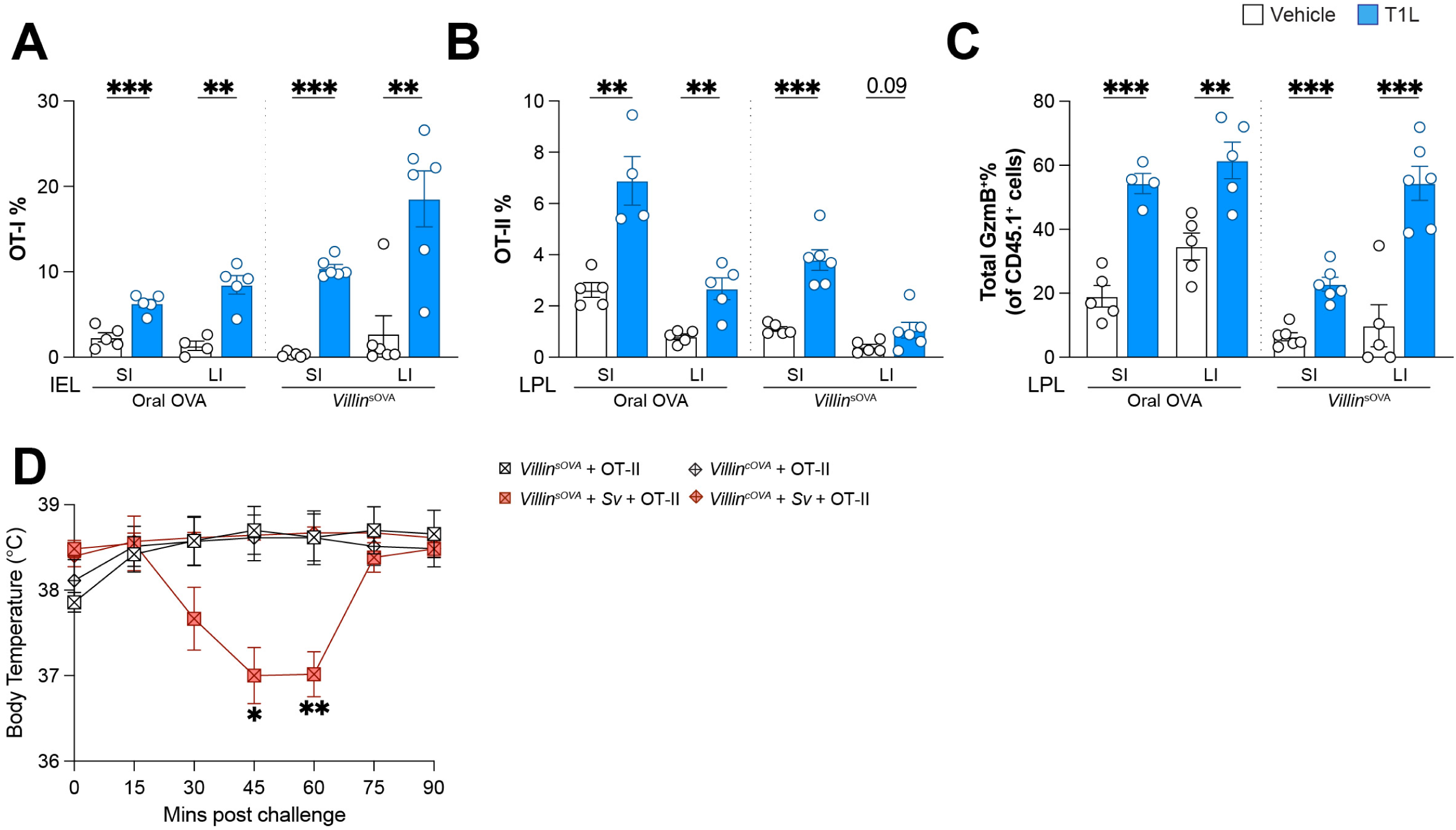
Intestinal viral infection leads to comparable pro-inflammatory T cell infiltration of the gut following oral and self-antigen recognition and helminth infection triggers breakdown in tolerance and anaphylaxis in response to secreted but not cytosolic OVA. A-C. Frequencies of OT-I among total CD8^+^ T cells (**A**) in the IEL fraction, OT-II among total CD4^+^ T cells (**B**) or GzmB^+^ OT-I among total OT-I cells (**C**) in the LPL fraction of the gut tissue of T1L-or vehicle-infected mice that were fed OVA or express sOVA (*n* = 4-6 per group, as indicated by symbols). Data are pooled from two independent experiments. **D**. Body temperature measurements of indicated groups every 15 min post 50 μg i.p. OVA challenge (*n* = 5-7 per group, as indicated by symbols). Data represent one experiment. * p<0.05, ** p<0.01, *** p<0.001 by t-test.

**Figure S7.**
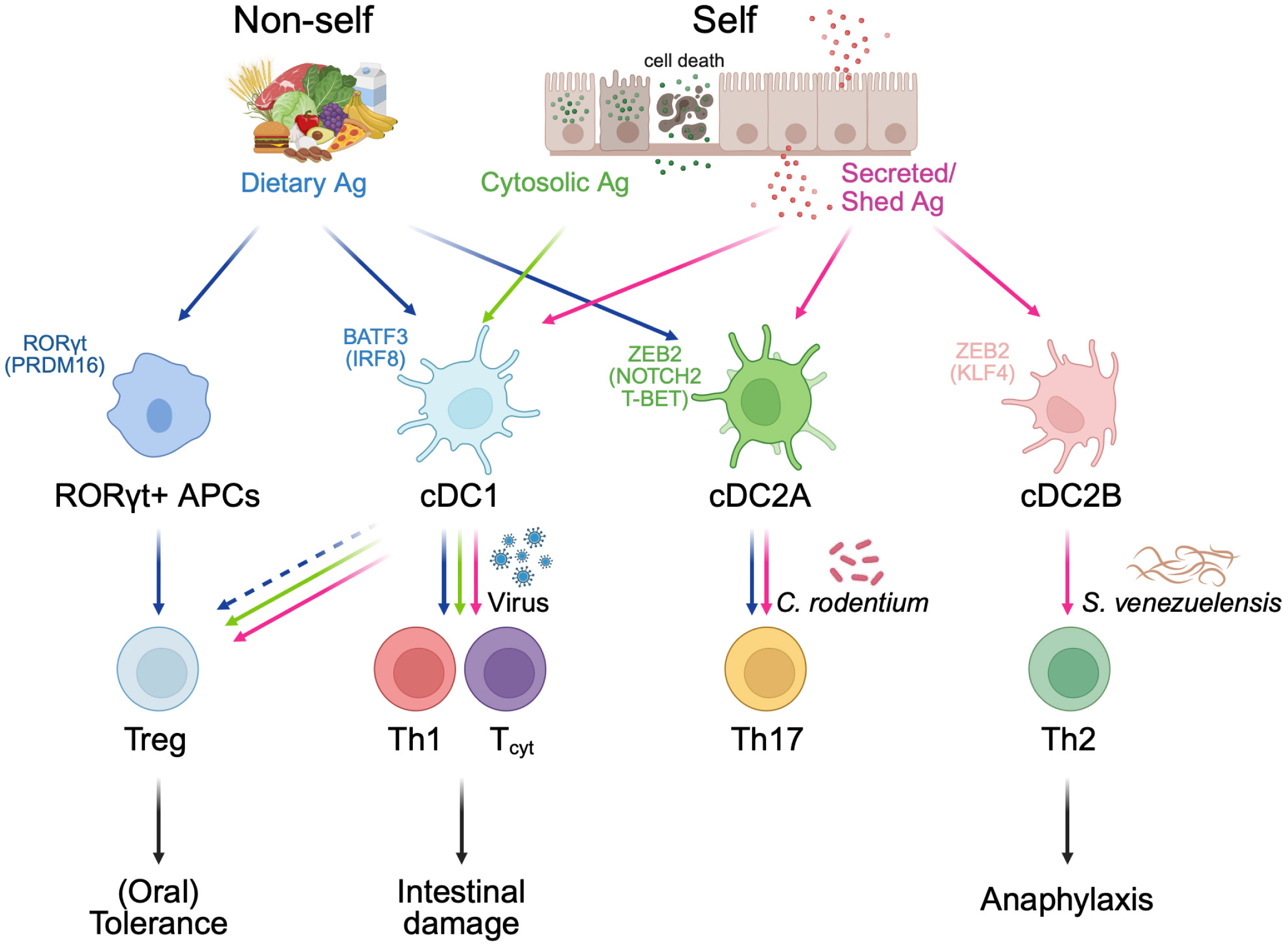
Graphical Abstract.

